# Initiation of Lumen Formation from Junctions via Differential Actomyosin Contractility Regulated by Dynamic Recruitment of Rasip1

**DOI:** 10.1101/2024.05.23.595279

**Authors:** Jianmin Yin, Niels Schellinx, Ludovico Maggi, Kathrin Gundel, Cora Wiesner, Maria Paraskevi Kotini, Minkyoung Lee, Li-Kun Phng, Heinz-Georg Belting, Markus Affolter

**Author notes:** Correspondence (J.Y.), (H.-G.B.), (M.A.).

## Abstract

Vascular lumens are essential for the dispersal of nutrients, oxygen, and immune cells throughout the body. *De novo* lumen formation necessitates the precise segregation of junctional proteins from apical surfaces. Direct visualization of this process and a comprehensive understanding on how the apical and junctional compartments are segregated and established remain elusive, due to the lack of visualizable systems, proper reporters and manipulative tools. Using zebrafish as an experimental model, we have established a wealth of molecular reporters, photo-convertible and optogenetic tools and combined them with genetic analyses to study the establishment of apical domains from pre-apical junctional complexes. Our study identified Rasip1 as one of the earliest apical proteins recruited, peripheralizing cortical contractility at the junctional patches by inhibiting NMII at the center, thereby initiating the outward flow of junctional complexes. After the establishment of the apical compartments, Rasip1 shuttles between junctions and the apical compartments to the high-tension sites and tunes down the local hyper contractility, restricting Cdh5 to junctions by suppressing apical contractility. Above all, the formation of *de novo* lumens and maintenance of junctional integrity depend on the precise control of contractility within the apical compartments and junctions, orchestrated by the dynamic recruitment of Rasip1.

## Introduction

Endothelial cells (ECs) polarize and generate apical domains and lumens to create a functional multicellular tubular network of blood vessels. Cell-cell contacts, formed by cadherin-based adherens junctions (AJs) and tight junctions, reside between the apical and basolateral membranes and seal the endothelial barrier. To establish appropriate *de novo* apical compartments, ECs must regulate proper segregation of junctional proteins from apical surfaces. However, the mechanisms governing the establishment and maintenance of distinct cellular compartments *in vivo* remain poorly understood. Obtaining a precise map of protein recruitment and segregation during apical polarization and a comprehensive understanding of this process had been proven to be extremely difficult *in vivo*, due to the lack of visualizable systems, proper reporters and manipulative tools.

The formation of a blood vessel network during embryogenesis requires both vasculogenesis and angiogenesis. Individual EC progenitors aggerate into cord structures that open central cavities during vasculogenesis (**Strilić et al., 2009; Xu & Cleaver, 2011**). Later during development, new vessels are primarily formed through angiogenesis, a process in which new vessels sprout from pre-existing ones, grow and connect to adjacent sprouts or other vessels (**Betz et al., 2016; Eelen et al., 2020; Schuermann et al., 2014**). Many cellular and molecular commonalities in forming *de novo* lumens have been revealed from studies in various contexts with different model organisms (**Datta et al., 2011; Sigurbjörnsdóttir et al., 2014**). In this study, we used the formation of the dorsal longitudinal anastomotic vessel (DLAV) in zebrafish as an experimental model to analyse the establishment of apical compartments *in vivo* (**Lawson & Weinstein, 2002**). During DLAV formation, a new vessel is built by the interaction of two approaching tip cells (**Betz et al., 2016**). Upon contact between the two tip cells, AJs proteins, such as Cdh5, are deposited at the contact site (Fig. 1A) (**Blum et al., 2008; Herwig et al., 2011**). At these adhesion sites, cells deposit *de novo* apical membranes while the initial adhesion spots open into junctional rings with apical membrane in between (Fig. 1A; Video 1) (**Betz et al., 2016**). Here, the key questions we aim to address are how different cellular compartments are established and maintained during *de novo* lumen formation.

**Figure 1.**
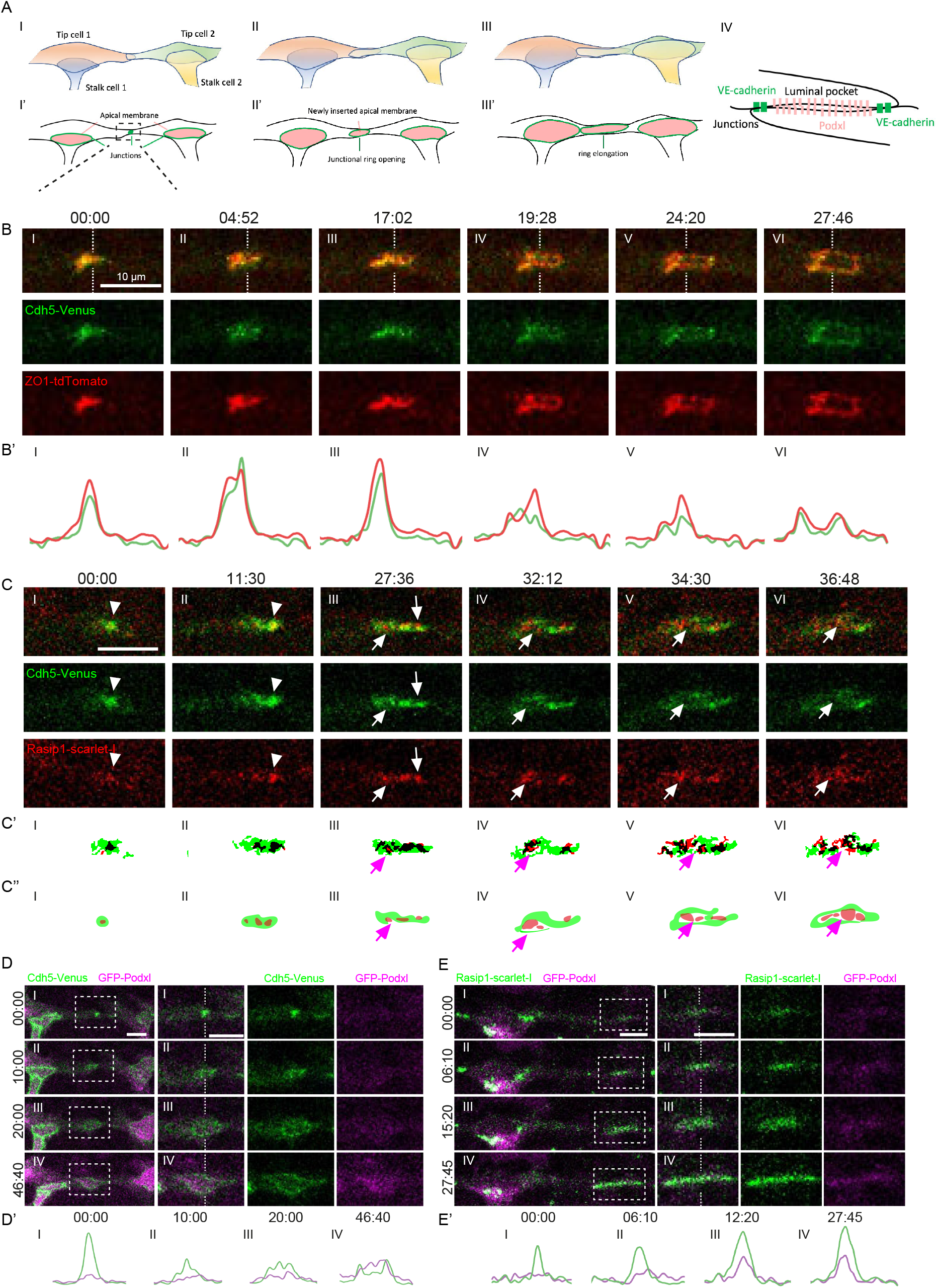
Rasip1 is involved in the patch-to-ring transition and establishment of apical compartments. (A)Schematic depicting of anastomosis of two tip cells in the formation of DLAV. Stable cell-cell contact site is formed with deposition of junctional proteins including Cdh5 (I). The cell-cell contact site opens into a ring structure while apical membrane is inserted into the luminal pocket within the junctional ring (II). Cell rearrangement leads to the elongation of junction ring (III). Transverse view of the luminal pocket located within junctional ring (IV). (B)Time lapse with expression of Cdh5-Venus and ZO1-tdTomato imaged from 30 hpf throughout the patch-ring transition. (B’) Intensities of Cdh5-Venus and ZO1-tdTomato along the dashed lines in (B). (C)Time lapses showing the recruitment of Rasip1 to the junctional patches and nascent junctional rings. White arrowheads label the Rasip1 clusters within the junctional patches. White arrows label the Rasip1 clusters localized at the periphery of junctional patches or within the nascent apical compartments. (C’) Automatic thresholding of Cdh5-Venus and Rasip1-scarlet-I in (C). Black color corresponds to the colocalized signals after thresholding. (C’’) Graphic diagram showing the localizations of signals in (C’). (D and E) Time lapses showing the recruitment of Podxl to the apical compartments together with Cdh5-Venus(D) and Rasip1-scarlet-I(E). (D’) Intensities of Cdh5-Venus and GFP-Podxl along the dashed lines in (D). (E’) Intensities of Rasip1-Scarlet-I and GFP-Podxl along the dashed lines in (E). All scale bars: 10 μm.

The actin cytoskeleton and its regulatory Rho GTPases are implicated in the EC polarization and junctional remodeling (**Beckers et al., 2010; Davis et al., 2011; Iruela-Arispe & Davis, 2009**). Rasip1 (Ras-interacting protein 1) is a potent regulator on Rho GTPase activity (**Barry et al., 2016; Post et al., 2013; Wilson et al., 2013; Xu et al., 2009**). Previous studies in mice and zebrafish suggest that Rasip1 is required for the clearance of apical junctions during vasculogenesis and angiogenesis. In mouse mutant embryos, the angioblasts fail to localize junctional proteins to the cord periphery during vasculogenesis of the dorsal aorta (DA) (**Koo et al., 2016; Xu et al., 2009, 2011**). Similarly, in zebrafish embryos undergoing DLAV formation, *rasip1* mutants manifest ectopic and reticulated junctional patterns within the presumed apical compartments (**Lee et al., 2021**).

Rasip1 has been shown to inhibit RhoA activity by binding to the GTPase-activating protein Arhgap29 (**Koo et al., 2016; Post et al., 2015; Xu et al., 2011**). A mechanism has been proposed that Rasip1 clears apical junctions via mechanical sliding of junctional proteins along actin tracks to the cord periphery through activating Cdc42 and NMII during vasculogenesis of the DA (**Barry et al., 2016**). However, direct visualization of this process, how Rasip1 possibly switches between activator and inhibitors of NMII, and how the apical and junctional contractilities are coordinated during *de novo* lumen formation remain elusive. To gain a comprehensive understanding of apical clearance, we explored both of the temporal and spatial dynamics of apical compartmentalization and junctional remodeling during *de novo* lumen formation. By developing and utilizing a wealth of molecular reporters, we identified Rasip1 as the earliest apical protein recruited to the initial cell-cell contact sites, segregating with Cdh5 during *de novo* lumen formation. Further, after the establishment of the apical compartment, Rasip1 shuttles between junctions and the apical compartments to the high-tension sites. Combined with advanced live imaging, photoconversion, pharmacological inhibition and the use of optogenetic tools, we demonstrated that Rasip1 initiates the outward flow of junctional complexes by and further stabilizes junctional complexes and constrains Cdh5 detachment by inhibiting apical contractility.

## Results

### Junctional remodeling and *de novo* lumen formation in the DLAV

Morphogenesis of the DLAV in the zebrafish is characterized by the de novo formation of a lumen between two neighboring sprouts. Here, luminal pockets form an apical compartment which later on connects to neighboring lumens to generate vascular patency (Fig. 1A). To elucidate the emergence of *de novo* apical compartments from initial cell-cell contact sites, we employed the Cdh5 reporter Cdh5-Venus [*Tg(cdh5: cdh5-TFP-TENS-Venus)*^*uq11bh*^] and the ZO-1 (tjp1a) reporter *TgKI(tjp1a-tdTomato)*^pd1224^ (**Lagendijk et al., 2017; Levic et al., 2021**). The two junctional markers strongly colocalized and exhibited a similar rearrangement pattern from junctional patches into junctional rings within an approximate 30-minute time window (Fig. 1B; Video 2). The initial cell-cell contact sites spread towards the periphery and formed junctional patches (Fig 1B(I-III)). These junctional patches voided inside and transited into nascent junctional rings (Fig. 1B(V)). Subsequently, the junctional rings further expanded, generating *de novo* apical compartments with minimal junctional materials inside (Fig. 1B(VI)).

We next explored the recruitment of apical proteins during the initial lumen formation of DLAV. Rasip1, a vascular-specific regulator of GTPases, has been implicated in the de novo formation of apical compartments during vasculogenesis and angiogenesis (**Barry et al., 2016; Lee et al., 2021**). We therefore generated a fluorescent reporter line [*Tg(fli1a: Rasip1-scarlet-I)*^*ubs59*^ to follow the distribution of Rasip during lumen formation in the DLAV. Consistent with antibody stainings, Rasip-scarlet-I was strongly recruited to the initial contract sites (Fig. 1C(I and II), S1A and S1B; Video 3). During the patch-ring transition, Rasip1 started to segregate from Cdh5 by translocating to the periphery as clusters (Fig. 1C(III and IV), S1A(I-III)). Further segregation of Rasip1 and Cdh5 resulted in spatial partitioning, with Cdh5 localized at the periphery and Rasip1 concentrated in the central region (Fig. 1C(IV-VI), S1A(IV and V)). After the patch-ring transition, most of the Rasip1 protein was found inside of the junctional rings at the presumed apical domains (Fig. 1C(VI), S1A(V), and S1C). In contrast, the recruitment of the surface glycoprotein Podocalyxin (PODXL) only became apparent in the nascent junctional rings after the patch-ring transition, as visualized by the Podxl1 live reporter [*Tg(fli1a: GFP-Podxl1)*^*ncv530Tg*^] and antibody staining (Fig. 1D (IV), S1D and S1E). The direct comparison between Rasip1 and Podxl1 further demonstrates that during DLAV anastomosis Rasip1 is recruited to the prospective apical membrane prior to Podocalyxin (Fig. 1E).

### Improper compartmentalization in *rasip1* mutants

Rasip1 has been implicated in apical junction clearance (**Barry et al., 2016; Lee et al., 2021**). To explore the temporal and spatial junctional dynamics of this process in both wild-type embryos and *rasip1* mutants, we utilized the *rasip1*^*ubs28*^ allele, which harbors a 35 kb deletion from exon 3 to 16(**Lee et al., 2021**). We first examined the shape of junctions via antibody staining of ZO-1 and Cdh5 at the DLAV. In wild-type embryos, junctions exhibited distinct ring-like structures characterized by well-defined boundaries and predominantly clean apical compartments, with limited presence of junctional proteins (Fig. S2A). In contrast, in *rasip1* mutants, ectopic clusters (white arrowhead) or linear structures (white arrow) of Cdh5 and ZO-1 appeared within the presumed apical compartments, thus designated as reticulated junctions (Fig. S2B). The boundaries of the presumed apical compartments appeared fuzzy and discontinuous, with widespread distribution of ZO-1 and Cdh5 (blue arrowhead) (Fig. S2B). We observed similar phenotypes with the Cdh5-Venus live reporter in *rasip1* mutants (Fig. 2A and 2B). By measuring the average levels of Cdh5-Venus within the boundary areas (green region in Fig. 2A and 2B) compared to apical compartments (red region in Fig. 2A and 2B), we confirmed the ectopic presence of junctional proteins within the apical compartments in *rasip1* mutants (Fig. 2C).

**Figure 2.**
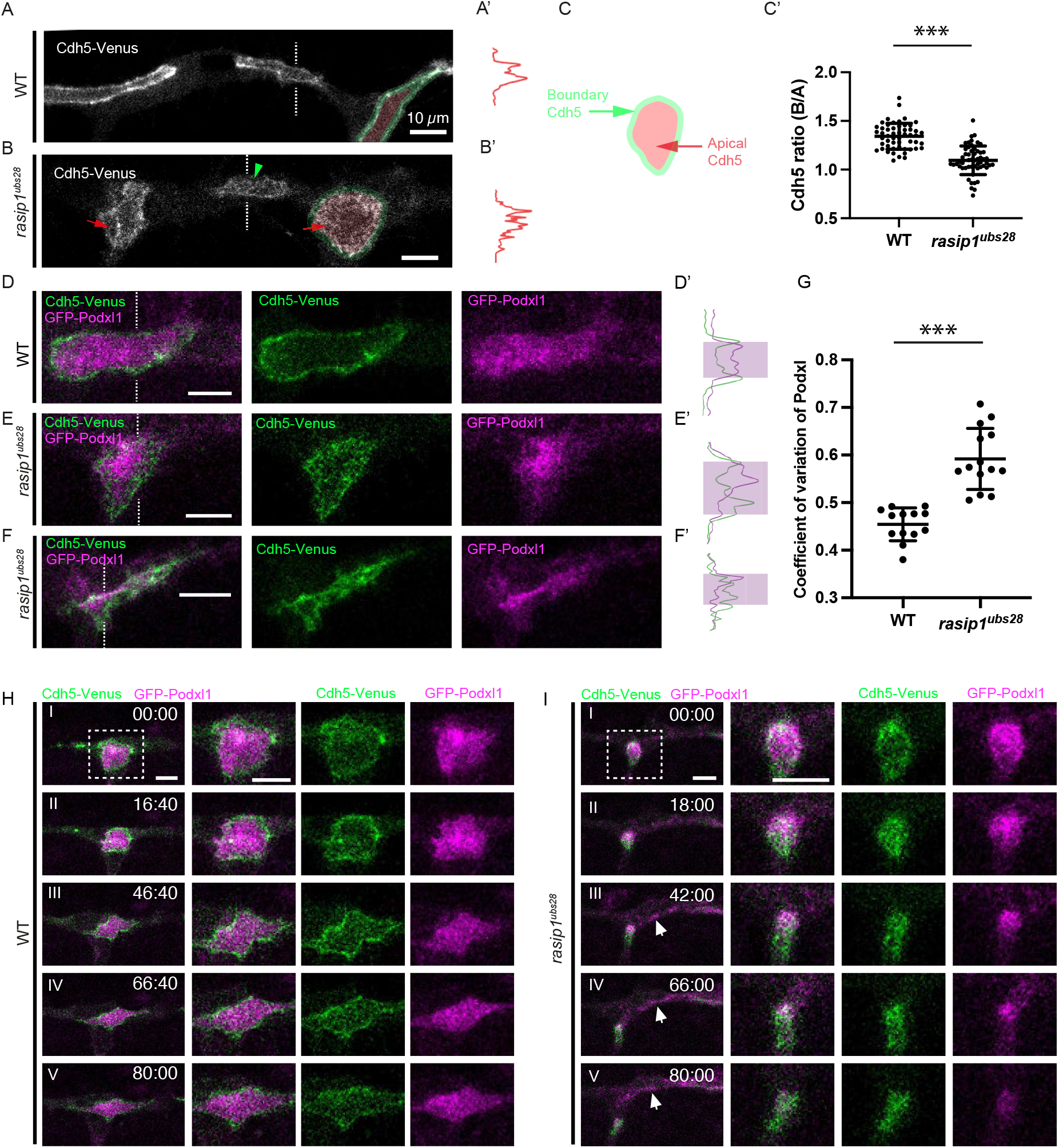
Apical clearance and lumenal defects in *rasip1* mutants. (A and B) Expression of Cdh5-Venus in wild-type embryos (A) and *rasip1* mutants (B) at 32 hpf. Red arrows label the junctional clusters in the presumed apical compartments in *rasip1* mutants. Green arrowhead labels the discontinuous junctions. (A’ and B’) Intensities of Cdh5-Venus along the dashed lines in (A) and (B). (C) Quantification of the ratio of the average Cdh5 intensity between the boundary regions (green masks in (A) and (B)) and the apical compartments (red masks in (A) and (B)) (WT: n=53; *rasip1*: n=55). (D-F) Cdh5-Venus and GFP-Podxl1 in wild-type embryos (D) and *rasip1* mutants (E and F). (D’–F’) Intensities of Cdh5-Venus and GFP-Podxl1 along the dashed lines in (D-F). (G) Coefficient of variation of Podxl1 per apical compartment with mean ± SD (WT: n=13; *rasip1*: n=14). (H and I) Time lapses of Cdh5-Venus and GFP-Podxl1 in wild-type embryos (H) and *rasip1* mutants (I). White arrows in (I) labels the GFP-Podxl1 emerged outside of junctions. All scale bars: 10 μm. *** P<0.001 (t-test).

Since loss of Rasip1 prevents the apical clearance of junctional proteins, we wanted to further explore the role of Rasip1 in the apical compartmentalization. We imaged GFP-Podxl1 to explore *de novo* lumen formation in both wild-type embryos and *rasip1* mutants in the DLAV. In wild-type embryos, Podxl1 displayed a largely uniform distribution inside the junctional rings (Fig. 2D and 2D’). In contrast, Podxl1 exhibited non-uniform aggregations within the reticulated junctions, with intensified signals concentrated in the center or the periphery of the apical compartments in *rasip1* mutants (Fig. 2E, 2E’, 2F and 2F’). Additional analyses revealed a significantly higher coefficient of variation of GFP-Podxl1 signal within the apical compartments in *rasip1* mutants (Fig. 2G). Live imaging of Cdh5-Venus and GFP-Podxl1 indicated that Podxl1 was restricted within the presumed apical compartments during the expansion of junctional rings in wild-type embryos (Fig. 2H). In contrast, ectopic Cdh5 appeared in the presumed apical compartments in *rasip1* mutants, while the Podxl1 signal was consistently lost within the apical compartments and became evident outside (Fig. 2I; Video4). These data show that Cdh5 and Podxl1 are less restricted in *rasip1* mutants, leading to improper compartmentalization of junctional and apical compartments.

### Insufficient restriction on junctional Cdh5 in r*asip1* mutants

Luminal defects in vasculogenesis in *rasip1* mutants have been proposed to be caused by the failure to remove junctional proteins from the apical membrane (**Barry et al., 2016; Lee et al., 2021**). To elucidate the mechanisms underlying the mislocalized junctional complexes in zebrafish DLAV, we created a photoconvertible Cdh5 reporter, Cdh5-mClav [*Tg(cdh5: cdh5-mClavGR2)*^*ubs58*^] (Fig. 3A). Different pools of Cdh5 were labelled by photo-conversions of Cdh5-mClav in discrete regions of interest (ROIs) at desired stages. We first selectively photoconverted entire junctional patches as indicated by the squares drawn around them (Fig. 3B). Upon complete conversion, we monitored subsequent junctional remodeling in both wild-type embryos and *rasip1* mutants (Fig. 3C and 3D). The photoconverted Cdh5 from the junctional patches rearranged towards the periphery and eventually became part of junctional rings together with newly incorporated green Cdh5 in both wild-type and *rasip1* mutant embryos (Fig. 3C(III) and 3D(III)). Converted Cdh5 slowly faded during subsequent junctional remodeling, possibly due to recycling and photo-bleaching in both conditions. Interestingly, in sharp contrast to wild-type embryos displaying elongating junctions, the junctional ring in *rasip1* mutants quickly transformed into a reticulated pattern, losing clear boundaries and displaying ectopic junctional complexes within the presumed apical domain (Fig. 3D(V)). Further time-lapse analyses on *rasip1* mutants revealed a large portion of the initial adhesion sites transitioned into ring shape temporally (24/35) but smeared later (22/24) (Fig. S3). Thus, we hypothesize that Rasip1 stabilizes junctional materials at the boundaries of the apical compartments.

**Figure 3.**
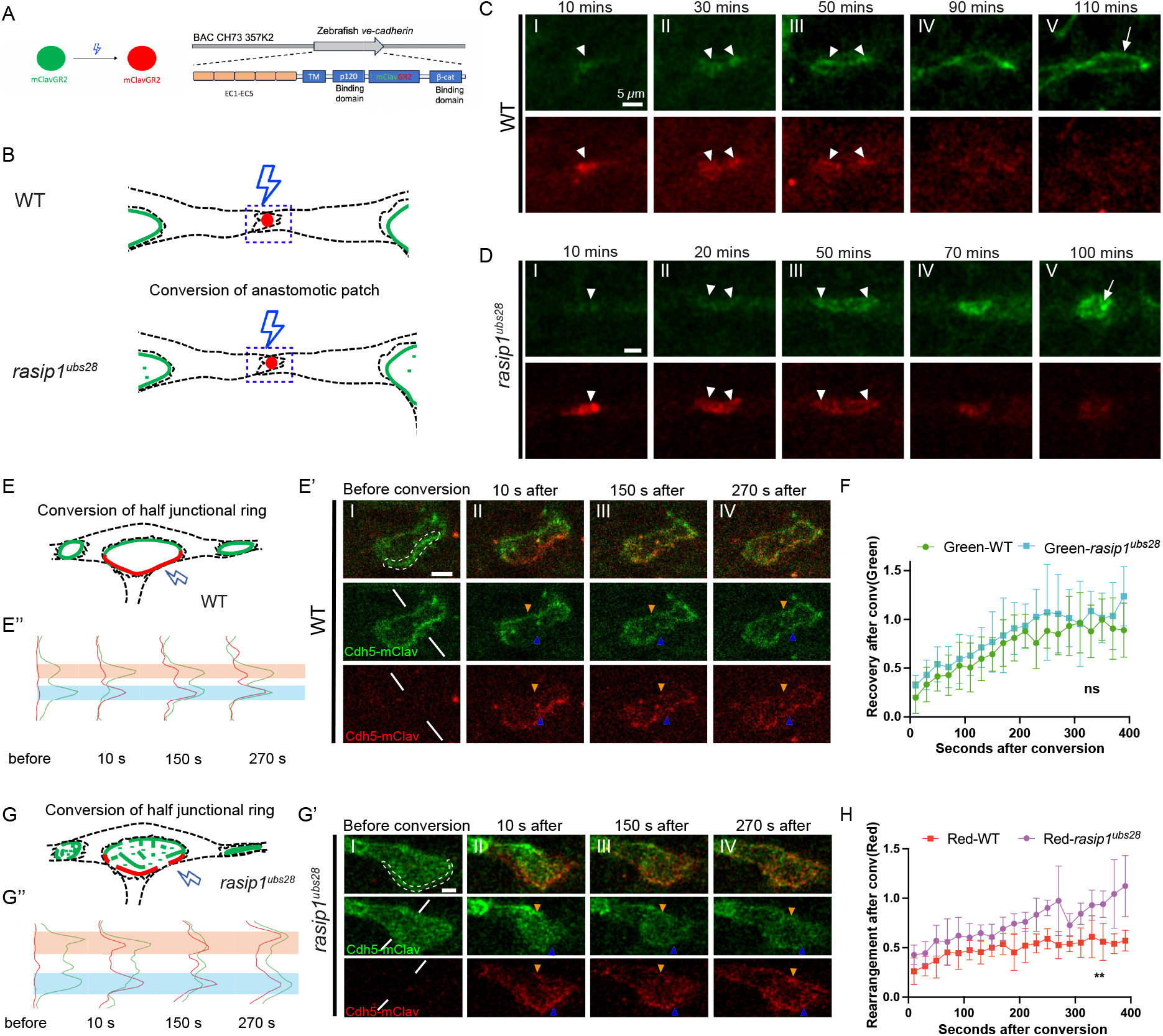
Cdh5 is poorly restricted to the junctions in *rasip1* mutants. (A)Schematic drawing of photo-conversion of mClavGR2 and *cdh5-mClav* cDNA recombined into cdh5 BAC clone. (B)Diagrams of photo-conversions at the initial cell-cell contact sites (anastomotic patches) in wild-type embryos and in *rasip1* mutants. (C and D) Time lapses of Cdh5-mClav after photo-conversion specifically at junctional patches. White arrowheads label the converted Cdh5-mClav. White arrow labels the reticulated junctions formed in *rasip1* mutants in D (IV). (E-H) Photo-conversions on half junctional rings in wild-type embryos (E) and equivalent regions in *rasip1* mutants (G). (E’ and G’) Time lapses of Cdh5-mClav after photo-conversion of half junctional rings in wild-type embryos (E’) and equivalent regions in *rasip1* mutants (G’). White dashed lines label converted region before conversion. Blue and orange arrowheads label the converted and unconverted half rings respectively. (E’’ and G’’) Intensities of green and red Cdh5-mClav along the white lines in (E’) and (G’). Blue and orange masks label the converted and unconverted half rings. (F) The relative intensity of green Cdh5-mClav on the converted half rings compared to the unconverted half rings in wild-type embryos (n=5) and *rasip1* mutants (n=6) with mean ± SD. (H) The relative intensity of red Cdh5-mClav on the unconverted half rings compared to the converted half rings in wild-type embryos (n=5) and *rasip1* mutants (n=6) with mean ± SD. All scale bars: 5 μm. ** P<0.01, ns P>0.05 (t-test on area under the curve).

To test this hypothesis, we selectively photo-converted half of the junction rings with hand drawn ROIs in wild-type embryos and in *rasip1* mutants (Fig. 3E-H; Video 5 and 6). Upon complete photo-conversion, all of the green Cdh5-mClav turned red in the converted region, whereas the other half of the ring remained unconverted (Fig. 3E’ and 3G’). We first quantified the recovery speed of green Cdh5-mClav in both wild-type and mutant embryos. In both conditions, the converted half rings took approximately 200 s to reach 90% of the green intensities of the unconverted half ring, suggesting a rapid incorporation of new junctional proteins in both conditions (Fig. 3F). Conversely, we also assessed to what extend red (converted) Cdh5 moved away from the converted region. In wild-type embryos, the converted half ring exhibited a red Cdh5 level two times higher than that of the unconverted half ring even after 400 s (Fig. 3E’(IV), 3E’’ and 3H). In contrast, a large amount of red Cdh5 appeared in the apical domain as clusters and in the unconverted half ring in *rasip1* mutants (Fig. 3G’(II-IV), 3G’’). The unconverted half ring displayed a comparable level of red Cdh5-mClav approximately 200-300 s after conversion (Fig. 3H). Thus, in *rasip1* mutants, junctional Cdh5 is poorly restricted, indicating that Rasip1 is required for the stabilization of junctions.

### Increased motility of Cdh5 clusters with inadequate reintegration in r*asip1* mutants

The above observations suggest that Rasip1 is required to stabilize Cdh5 at cell-cell junctions. In the absence of Rasip1, Cdh5 exhibited increased motility towards the apical compartment. In order to better understand Cdh5 motility, we tracked individual junctional particles in the apical compartment. In wild-type embryos, a very small number of Cdh5-Venus particles appeared within the apical compartments (Fig. 4A, red arrowhead; Video 7). In contrast, a larger number of clusters or linear fragments of Cdh5 detached from the boundaries and moved towards the apical domain in *rasip1* mutants (Fig. 4B and 4C, red arrowhead; Video 8 and 9). Interestingly, new Cdh5 was deposited at the boundaries, while pieces of old junctional clusters or fragments thereof detached (Fig. 4B and 4C, green arrowhead). The detachment of junctional clusters in *rasip1* mutants was further confirmed by observing the movement of photo-converted particles from junctions into the apical domains (Fig. S4A).

**Figure 4.**
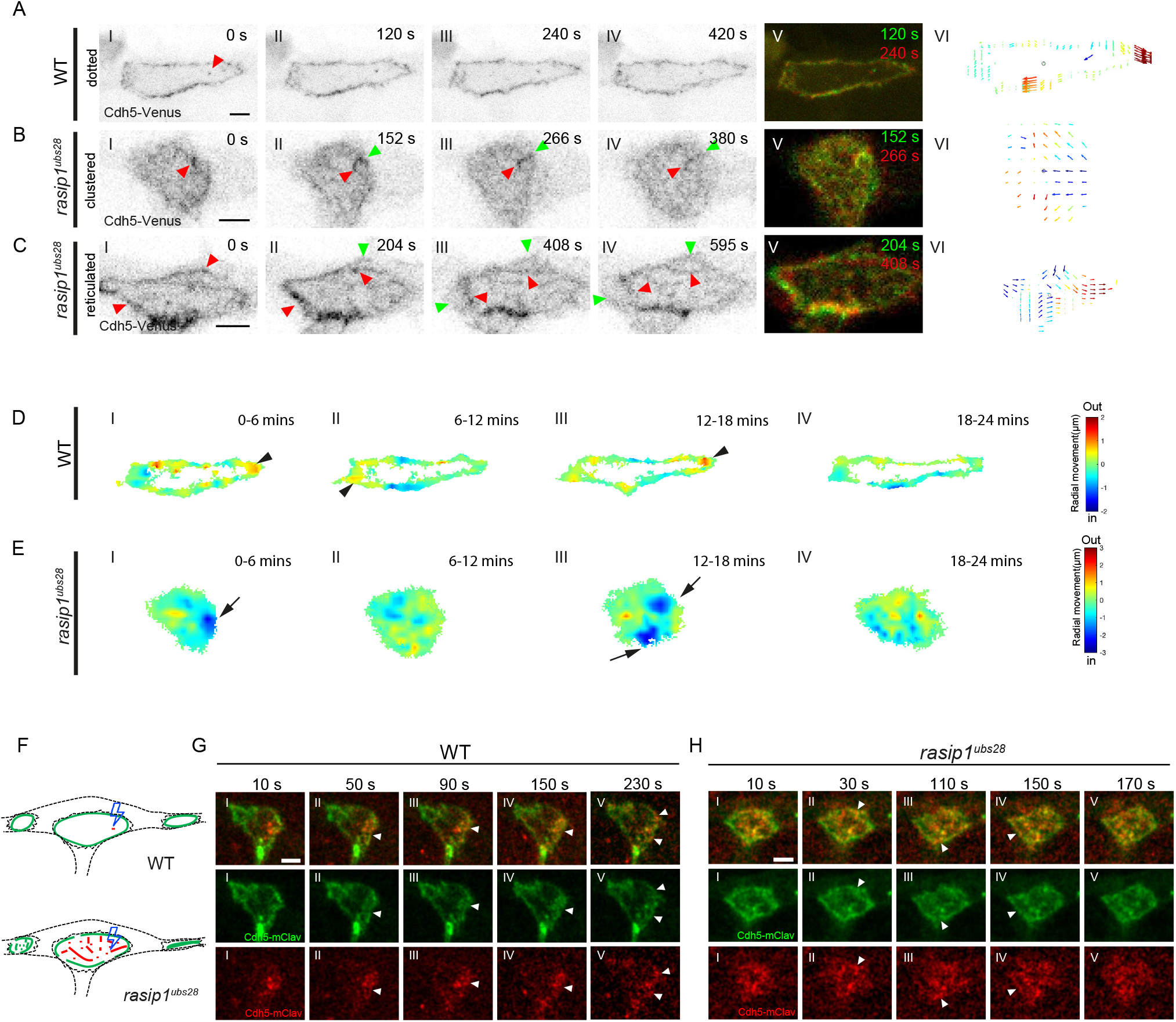
Cdh5 clusters detach from junctions and move towards the apical compartments in *rasip1*^*ubs28*^ mutants. (A-C) Time lapses of Cdh5-Venus in WT embryos (A) and in *rasip1*^*ubs28*^ mutants (B and C). Red arrowheads label the dots, clusters or linear structures of Cdh5 within the apical compartments. Green arrowheads label the newly formed junctions at the boundaries after the detachment of old junctions. (VI) Arrow maps showing the speed and directions of the flow of Cdh5 between the two time points in (V). The quivers were colored according to the radial velocity relative to the centers of the apical compartments. (D and E) Heat maps showing the averaged velocity of Cdh5 at the radial direction at 6 minutes interval in WT embryos (D) and *rasip1*^*ubs28*^ mutants (E). Averaged velocity maps were generated from two frames with a 36-second interval using PIV and were further combined into a heat map displaying the total particle movement over 360 seconds in the radial direction. Black arrowheads label outward movement while black arrows label inward movement. (F-H) Diagrams and timelapses of photo-conversion on the junctional clusters in the apical compartments of wild-type embryos (G) and *rasip1* mutants (H). White arrowheads label the converted Cdh5-mClav that moved to the boundaries. All scale bars: 5 μm.

To gain a better understanding on junction dynamics, we performed Particle Image Velocimetry (PIV) on high temporospatial resolution movies, enabling us to track and quantify Cdh5 movements both in the apical membrane and at the junctional boundaries (Fig. 4A(VI)-4C(VI)) (**Thielicke & Sonntag, 2021**). Significant inward movements toward the apical compartments were detected in *rasip1* mutants at both the boundary regions and in the apical compartments. Averaged velocity maps were generated at 6-minute intervals along the radial direction (Fig. 4D and 4E). In wild-type embryos, most of the movements occurred at the lateral side of junctional rings towards outside (red), correlating well with the lateral expansion of junctional rings (Fig. 4D). In contrast, significant inward (blue) movements of Cdh5 towards the center were observed in intermittent subregions in *rasip1* mutants (Fig. 4E). A closer examination of the velocity map revealed multiple types of apical Cdh5 movements in *rasip1* mutants: apical Cdh5 either dispersed, aggregated within the apical compartments, detached from boundaries, or attached to them (Fig. S4B; Video 10).

To further explore the fate of apical Cdh5, we selectively photo-converted the junctional particles in both conditions (Fig. 4F). In wild-type embryos, the apical Cdh5 particles dispersed, moved towards the boundaries and were incorporated into junctional rings (Fig. 4G). Similar incorporation of Cdh5 from the apical compartments to the boundaries was also observed in *rasip1* mutants, suggesting that the movements of Cdh5 clusters are bidirectional in these mutants (Fig. 4H). However, the converted Cdh5 that was incorporated into the boundaries appeared transient at junctions in *rasip1* mutants (Fig. 4H(II-IV), white arrowhead). Overall, our study indicates that loss of Rasip1 leads to increased motility of Cdh5 from junctions to the apical compartments and ineffective incorporation from the apical compartments back into cell-cell junctions.

### Precise control of actomyosin contractility is required for junctional remodeling and *de novo* lumen formation

Rasip1 has been proposed to inhibit RhoA or activate Rac1 and Cdc42, thereby either inhibiting or activating actomyosin contractility (**Barry et al., 2016; Post et al., 2015; Xu et al., 2011**). We therefore reasoned that some of the defects in junctional dynamics, which we observe in *rasip1* mutants may be caused by a dysregulation of actomyosin contractility. To explore the temporal-spatial activities of actomyosin contractility during *de novo* lumen formation and distinguish between the diverse roles of Rasip1 in this process, we utilized a NMII reporter, Myl9a-GFP (myosin light chain 9a-GFP) [*Tg(kdrl: Myl9a-GFP)*^*ip5Tg*^] (**Lancino et al., 2018**). In wild-type embryos, Myl9a largely colocalized with the junctional patches before the patch-to-ring transition (Fig. 5A(I and II); Video 11). However, Myl9a clusters emerged at the periphery and were devoid from the center of junctional patches prior to the ring formation (Fig. 5A(III)). After the nascent junctional rings were formed, Myl9a largely localized along the junctions (Fig. 5A(IV)). Similar transitions in the localization of Myosin were confirmed with the Myl9b-Cherry reporter we created (myosin light chain 9b-mCherry) [*Tg(fli:gal4; UAS: Myl9b-mCherry)*^*rk32*^] (Fig. S5). Thus, relocalization of Myl9 precedes the formation of a defined junctional ring. Live imaging of Myl9a and Rasip1 indicated that Rasip1 colocalized first with Myl9a at the junctional patches (Fig. 5B(I and II)) and segregated later, with Myl9a localized at the periphery and Rasip1 in the center (Fig. 5B(III)).

**Figure 5.**
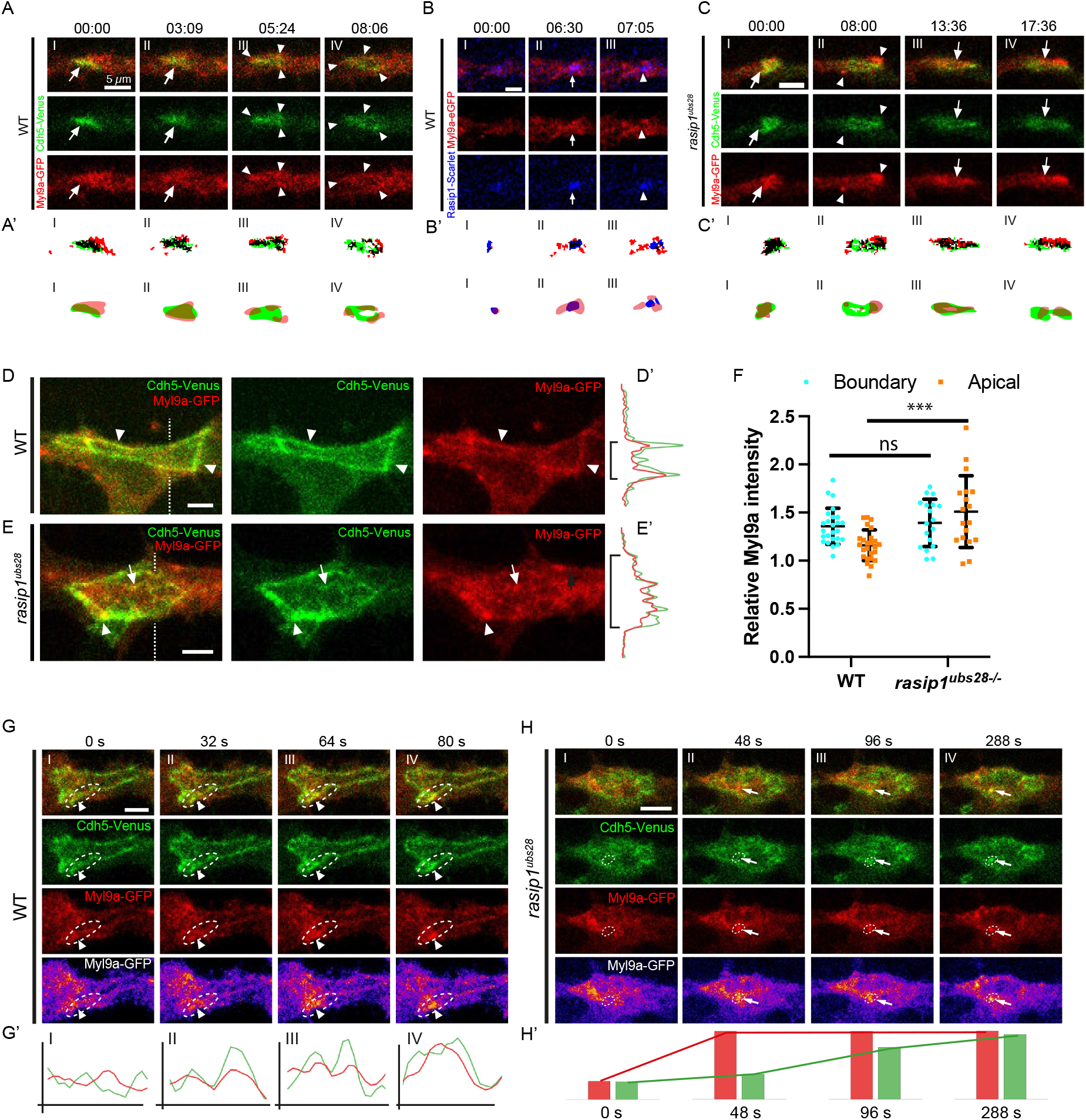
Rasip1 inhibits contractility at the junctional patches and apical compartments and determines how junctions remodel. (A-C) Cdh5-Venus and Myl9a-GFP in wild-type embryos (A), Rasip1-scarlet-I and Myl9a-GFP in wild-type embryos (B) and Cdh5-Venus and Myl9a-GFP in *rasip1* mutants (C) throughout the patch-to-ring transitions. White arrows label colocalization while white arrowheads label segregation of corresponding signals. (A’-C’) Automatic thresholding of corresponding signals in (A-C) and corresponding graphic diagrams. Black color corresponds to the colocalized signals after thresholding. (D and E) Distinct localizations of Myl9a-GFP in wild-type embryos (D) and *rasip1* mutants (E). White arrowheads label Myl9a at junctions. White arrows label Myl9a clusters within the presumed apical compartments in *rasip1* mutants. (D’ and E’) Intensities of Cdh5-Venus and Myl9a-GFP along the dashed lines in (D) and (E). (F) Normalized Myl9a intensities at the peripheral regions and the apical compartments in wild-type embryos and *rasip1* mutants (WT: n=25; *rasip1*: n=18) with mean ± SD. The Myl9a-GFP intensity is standardized using the mean signal level from the entire cell. (G and H) Time lapses of Cdh5-Venus and Myl9a-GFP showing synchronized clustering of Myl9a and Cdh5 in WT embryos (G) and in *rasip1* mutants (H). White arrowheads label the Cdh5 and Myl9a along the junctions (G). White arrows label the Cdh5 and Myl9a clusters within the presumed apical compartment (H). (G’) White arrowheads label the Cdh5 and Myl9a along the junctions marked in (G). (H’) Changes of the average signal levels of Myl9a-GFP and Cdh5-Venus within the marked area in (H). All scale bars: 5 μm. *** P<0.001, ns P>0.05(t-test).

In *rasip1* mutants, Myl9a was initially enriched in the junctional patches and subsequently appeared at the periphery, similar to wild-type embryos (Fig. 5C(I and II); Video 12). However, Myl9a soon lost its peripheral localization and emerged again in the junctional patches in *rasip1* mutants (Fig. 5C(III and IV)). We next examined the localizations of Myl9a-GFP at junctional rings and reticulated junctions in wild-type embryos and *rasip1* mutants (Fig. 5D and 5E). In wild-type embryos, most of the Myl9a-GFP localized along the junctions (Fig. 5D and D’). By contrast, in *rasip1* mutants, Myl9a predominantly localized at the apical domains as clusters (Fig. 5E and E’). Quantifications of the fluorescent intensities indicated that *rasip1* mutants displayed a higher overall level of Myl9a at the apical compartments compared to wild-type embryos (Fig. 5F). Although the Myl9a signals at the boundaries remained largely at the same level between wild-type and *rasip1* mutants, they exhibited distinct spatial arrangements. In wild-type embryos, Myl9a localized linearly along the junctions whereas in *rasip1* mutants, it exhibited intermittent clustering along the border (Fig. 5D-F).

The cortical actomyosin machinery transmits force to cell−cell contacts, thereby regulating junctional remodeling and cell shape. High-resolution imaging of Myl9a-GFP revealed the dynamics of Myl9a during junctional remodeling. In wild-type embryos, the Myl9a signals enriched along the junctions into local hotspots with aggregation of Cdh5 (Fig. 5G and 5G’; Video 13). In contrast, Myl9a predominantly aggregated within the apical compartments in *rasip1* mutants (Fig. 5H; Video 14). During this process, new Cdh5 clusters formed within the apical compartments, exhibiting a strong temporal-spatial correlation with the ectopic apical Myl9a clustering in *rasip1* mutants (Fig. 5H and 5H’).

### Rasip1 shuttles between apical and junctional compartments to the high contractility sites

To further explore the possible roles of Rasip1 in regulating contractility at junctions and within the apical compartment, we expressed C-terminal tagged Rasip1-Scarlet-I, as mentioned above, or N-terminal tagged GFP-Rasip1 in the zebrafish vasculature verified their functions by rescuing the apical clearance defects (Fig. 6A, S6A and S6B). Consistent with antibody staining, GFP-Rasip1 and Rasip1-Scarlet-I were highly confined within the junctional rings (Fig. 6B and 6C). Yet, Rasip1 distribution within the apical membrane exhibited non-uniformity, with clustered or linear arrangements proximal to junctions, as evidenced by antibody staining and Rasip1 reporter analyses (Fig. 6B and 6C, S6C). Live imaging of Rasip1 reporters revealed that Rasip1 was highly enriched at the local constricting sites of junctions and the apical compartments (Fig. 6D and 6E; Video 15). Apical Rasip1 translocated to the constricting site of junctions from the apical compartment (Fig. 6F(II); Video 16). After local contractions, the junctional Rasip1 translocated back to the apical domains (Fig. 6D(IV) and 6G; Video 17). These observations suggested that Rasip1 shuttled between the apical and junctional compartments. Live imaging of Rasip1 and Podxl1 showed that Podxl1 was not dynamically shuttled from the apical compartments to junctions until the very end of constriction, suggesting the dynamic translocation of Rasip1 is not due to the local aggregations of apical membrane (Fig. 6H).

**Figure 6.**
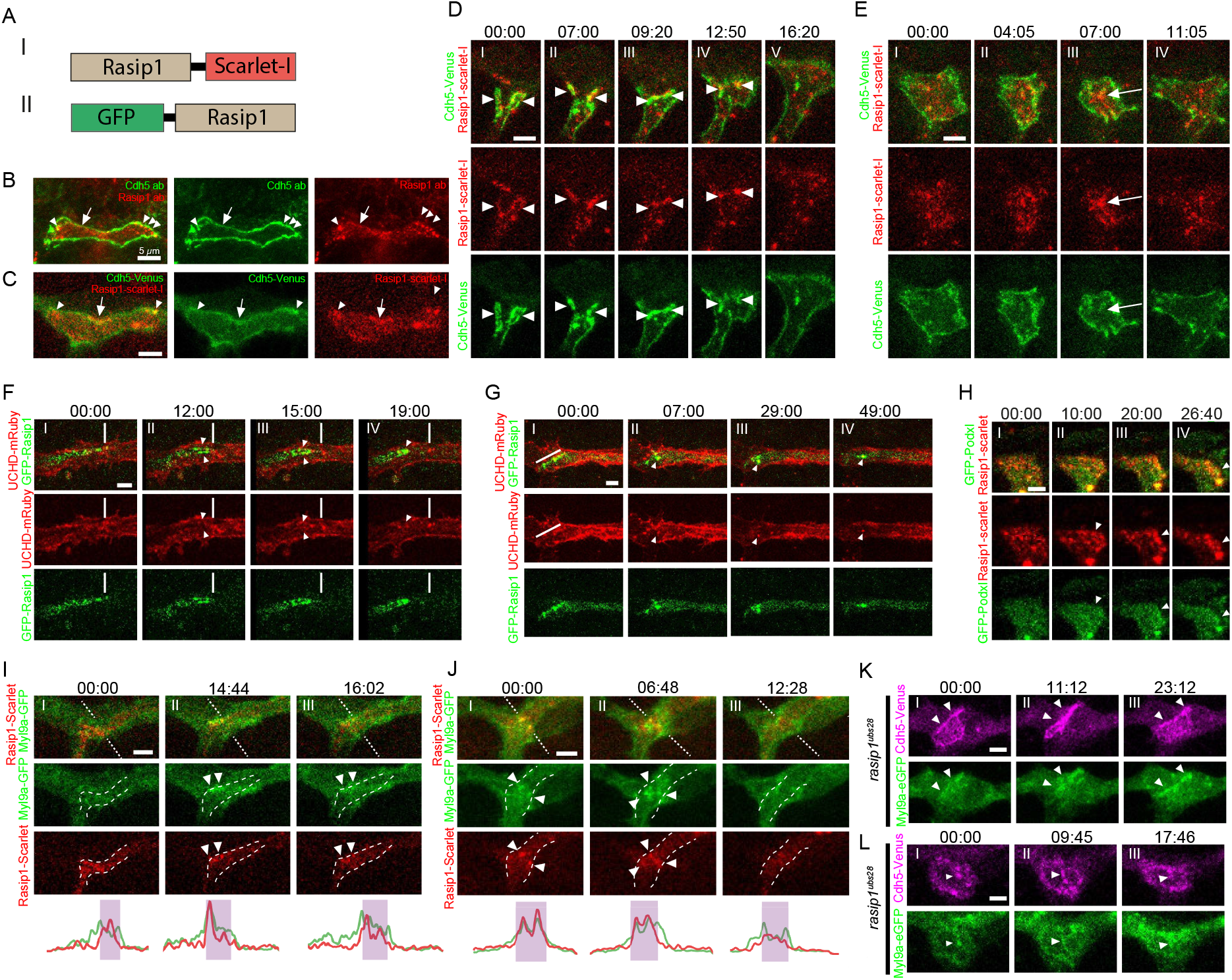
Rasip1 shuttles between junctions and the apical compartments in close association with Myosin. (A)Schematic drawing of recombined Rasip1-scarlet-I and GFP-Rasip1. (B)Antibody staining of Rasip1 and Cdh5 in wild-type embryos at 32 hpf. (C)Expression of Rasip1-scarlet-I and Cdh5-Venus in wild-type embryos at 32 hpf. For (B) and (C), white arrowheads label Rasip1 clusters whilst white arrows label linear Rasip1 localizations along the junctions. (D)Time lapses of Cdh5-Venus and Rasip1-scarelt-I with Rasip1 enriched at constricting junctions (white arrowheads). (E)Time lapses of Cdh5-Venus and Rasip1-scarelt-I with Rasip1 enriched at the apical compartment (white arrow). (F)GFP-Rasip1 relocated from the apical compartment to the constricting junctions. White lines label the position of the apical compartment before contraction. White arrowheads label the GFP-Rasip1 enriched at the constricting junctions. (G)GFP-Rasip1 relocated from junctions to the apical compartments as clusters. White line labels GFP-Rasip1 localized close to the anterior boundary of the apical compartment. White arrowheads label the GFP-Rasip1 clusters within the apical compartment. (H)Time lapses of GFP-Podxl1 and Rasip1-scarelt-I during constrictions. White arrowheads label the highly enriched Rasip1 at the constricting sites. (I and J) Time lapses of local enrichment of Myl9a-GFP and Rasip1-scarlet-I and subsequent depletion along the junctions (I) or within the apical compartments (J). White arrowheads label colocalized Rasip1 and Myl9a clusters. White dashed lines label boundaries of the apical compartments. (I’ and J’) Intensities of Myl9a-GFP and Rasip1-Scarlet along the dashed lines in (I) and (J). (K and L) Persisted excessive enrichment of Myl9a-GFP at junctions (K) or within the apical compartments (L) in *rasip1*^*ubs28*^ mutants labelled with white arrowheads. All scale bars: 5 μm.

Similar to the segregation observed in junctional patches, Rasip1 colocalized first and segregated later with Myl9a at junctions and within the apical compartments. Rasip1 was strongly enriched in clusters together with Myl9a hotspots at junctions or within the apical compartments (Fig. 6I(II) and 6J(I and II), Video 18 and 19). Both of Rasip1 and Myl9a clusters diminished later after their coexistence at the Myl9a hotspots (Fig. 6I(III) and 6J(III)). Thus, Rasip1 appears to dynamically respond to the local high contractility at junctions and the apical compartments. In combination with the functional studies on Rasip1, we hypothesized that Rasip1 inhibited contraction by localizing to the contracting sites either at the apical compartments or at the junctions. The presence of Rasip1 did not instantly halt contraction (Fig 6D and 6F). Instead, it appeared crucial in preventing excessive activation of the contractility machinery. In line with this hypothesis, in *rasip1* mutants, we observed sustained Myl9a clustering at junctions and within the apical compartments (Fig. 6K and 6L).

### Apical clearance defects can be restored or replicated through the modulation of apical actomyosin contractility

The above observations indicate a local ectopic elevation of actomyosin contractility within the apical compartments in *rasip1* mutants. To explore whether the mislocalization of Cdh5 in *rasip1* mutants is attributable to increased actomyosin tension, we modulated actomyosin tension through acute pharmacological inhibition of ROCK using Y-27632. Strikingly, Rock inhibition was sufficient to rescue several aspects of the *rasip1* mutant phenotypes. The initial adhesion sites opened properly and maintained clear ring shapes 80 minutes post-treatment compared to control embryos (Fig. 7A and 7B). The reticulated junctions that had formed prior to drug treatment were also rescued: the previously fuzzy boundaries became more defined, the apical compartments were cleared of junctional proteins and the elevated apical Myl9a was reduced under acute inhibition of ROCK (Fig. 7C, 7D and S7A). In wild-type embryos, acute ROCK inhibition with Y-27632 did not significantly alter the morphologies of junctions (Fig. S7B).

**Figure 7.**
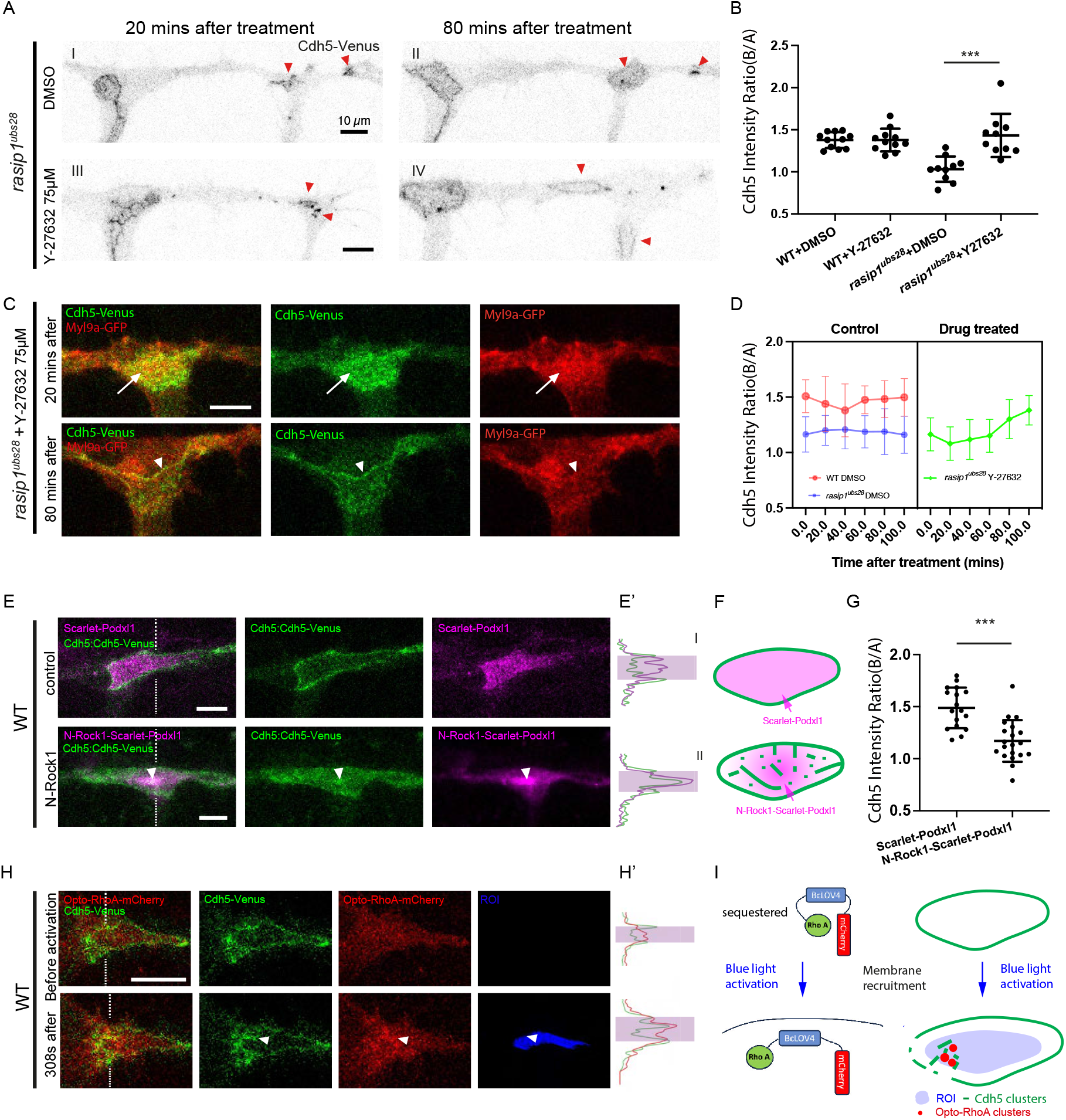
Precise control of apical contractility is required for apical clearance. (A)Acute DMSO or Y-27632 treatment on *rasip1* mutants with unopened cell-cell contact sites. (B)The Cdh5 boundary-to-apical ratio in the groups of WT+DMSO (n=11), WT+Y-27632(n=11), *rasip1* +DMSO(n=10), *rasip1* +Y-27632 (n=10) at 200 minutes after treatments for the newly opened junctions with mean ± SD. (C) Myl9a-GFP and Cdh5-Venus at 20 minutes and 80 minutes after acute Y-27632 treatment at 75 μM in *rasip1*^*ubs28*^ mutants. White arrows label the ectopic apical Myl9a and Cdh5 while white arrowheads label the junctional Myl9a at 80 minutes after Y-27632 treatment. (D) The Cdh5 boundary-to-apical ratio over the acute treatments in the groups of WT+DMSO (n=16), *rasip1* +DMSO (n=20), *rasip1* +Y-27632 (n=10) with mean ± SD. (E and F) wild-type embryos with the expression of Scarlet-I-Podxl1 or N-Rock1-Scarlet-I-Podxl (E) and corresponding diagram (F). (E’) Intensities of Cdh5-Venus and Scarlet-I-Podxl1/N-Rock1-Scarlet-I-Podxl along the dashed lines in (E). (G) Quantification of the Cdh5 boundary-to-apical ratio in wild-type embryos with the expression of Scarlet-I-Podxl1 (N=17) or N-Rock1-Scarlet-I-Podxl (N=20). (H and I) Activation of opto-RhoA selectively at the apical compartment (H) and corresponding diagram (I). White arrowhead labels the apical Cdh5 and opto-RhoA clusters. (H’) Intensities of Cdh5-Venus and opto-RhoA-mCherry along the dashed lines in (H). All scale bars: 10 μm. *** P<0.001, ns P>0.05(t-test).

To test whether the apical clearance defects in *rasip1* mutants can be phenocopied by elevating cortical actomyosin contractility specifically in the apical compartment, we overexpressed the catalytic domain of ROCK1 (N-Rock1) at the apical membrane by tagging it to the apically localized protein Podxl1. In agreement with this notion, Cdh5 ectopically localized within the apical compartments upon the expression of N-Rock1-Scarlet-Podxl1 (Fig. 7E-G). Unlike the evenly distributed Scarlet-Podxl1 observed in the apical membrane, N-Rock1-Scarlet-Podxl1 exhibited heterogeneous aggregations reminiscent of Podxl1 distribution in *rasip1* mutants (Fig 7E and 7F). To achieve precise temporospatial control of RhoA/Rock activities, we applied a single-component optogenetic tool for inducible RhoA GTPase signaling in zebrafish vasculature, *Tg(fli1a: RhoA-BcLOV4-mCherry)* (**Berlew et al., 2021**). Local RhoA signaling was activated under 450nm laser at hand drawn ROIs. After continuous 450nm laser activation within the apical compartments in wild-type embryos, Cdh5 clusters emerged at the apical compartment together with the RhoA-BcLOV4-mCherry clusters, while the boundary Cdh5 became fractured and fuzzy (Fig 7H and 7I; Video 20). In summary, these findings suggest that the increased apical contractility in *rasip1* mutants destabilizes the junctional complexes at the boundaries of the apical compartments.

## Discussion

### Rasip1 restricts Cdh5 to junctions by suppressing apical contractility

The removal of preapical adhesions is a key step in the formation of the apical compartment during *de novo* lumen formation. A previous study on vasculogenesis in mouse embryos proposed that junctional complexes were pulled from the preapical patch to the periphery by actomyosin contractility (**Barry et al., 2016**). Indeed, in this study, we observed Cdh5 rearrangement from the preapical patch to the periphery through photo-conversion of junctional patches. However, we also determined that it is not only the rearrangement, but also the proper restriction of junctional clusters that facilitates the “clearance” of junctional patches, as demonstrated through photo-conversion and particle tracking. VE-cadherin, the principal endothelial cell–cell adhesion molecule, responds to mechanical forces from the actomyosin cytoskeleton (**Dorland & Huveneers, 2017**). In wild-type embryos, most of the Myl9a and Myl9b localize along the junctions. In contrast, we detected elevated apical contractility in *rasip1* mutants, showing a strong correlation with the formation of apical Cdh5 clusters. Futhermore, we showed that the apical clearance defects in *rasip1* mutants can be rescued or phenocopied through pharmacological inhibition of ROCK or activation of RhoA at the apical compartments by opto-RhoA, indicating Rasip1 restricts Cdh5 to junctions by supressing apical contractility.

### Podxl1 and Rasip1 play distinct roles in lumen formation

The process of lumen formation during vasculogenesis and angiogenesis involves distinct stages (**Strilić et al., 2009**). Initially, endothelial cells establish cell-cell adhesion, followed by junctional relocalization, leading to the development of an apical interface between adjacent endothelial cells flanked by junctions. Subsequently, the incipient apical surfaces between two endothelial cells de-adhere from each other and create a small slit, initiating formation of lumenal space. Our investigation highlights the involvement of Rasip1 in precisely regulating junctional and apical contractility to facilitate the partitioning of junctions and apical domains. Conversely, a longstanding hypothesis suggests that the negatively charged sialic acids of Podxl1 and CD34 induce electrostatic repulsion, aiding in the separation of luminal surfaces at later stages (**Robbins & Beitel, 2010; Strilić et al., 2009**). Manipulations to neutralize sialic acid charge emphasize the indispensable role of Podxl1 in electrostatic repulsion during vascular lumen formation (**Strilić et al., 2010**). In our study, we identified that Rasip1 is initially recruited to junctional patches prior to the recruitment of Podxl1. This observation is consistent with the different roles of Rasip1 and Podxl1 at distinct phases of lumen formation. While Podxl1 is crucial for repelling apical surfaces between adjacent ECs, Rasip1 facilitates the proper segregation of apical membrane from junctions as a pre-step. In the absence of Rasip1, Podxl1 displays aberrant aggregations at the presumed apical comparments under enhanced apical contractility and becomes inadequately restricted to the apical domains.

### Defining cellular compartments by distinct levels of actomyosin contractility

One of the fundamental cell biological question is how distinct cellular compartment are established and maintained to achieve precise spatiotemporal regulation of numerous processes and pathways. Convincing evidence emerged from *in vitro* measurement on reconstructed model membranes showing that actomyosin contraction mediates macroscopic de-mixing of their membrane components and membrane compartmentalization (**Honigmann et al., 2014; Krapf, 2018; Vogel et al., 2017**). However, it remains largely unknow whether and how actomyosin contractility alters the segregation behavior of different cellular domains *in vivo*. In this study, we identified the roles of Rasip1 in the clear partitioning of the apical and junctional compartments by defining different levels of actomyosin contractility. According to our knowledge, Rasip1 is one of the earliest apical proteins recruited to the initial cell-cell contact sites (Fig. 8A(I)). Rasip1 and Myosin rearrange dynamically within the junctional patches and start to segregate with each other (Fig.8A(II and III)). In our model, the centrally localized Rasip1 inhibits Myosin, while Myosin localized at the periphery pulls the junctional complexes to borders and stabilizes them (Fig. 8B). In the absence of Rasip1, the junctional complexes are poorly restricted along the borders under increased apical contractility (Fig. 8C).

**Figure 8.**
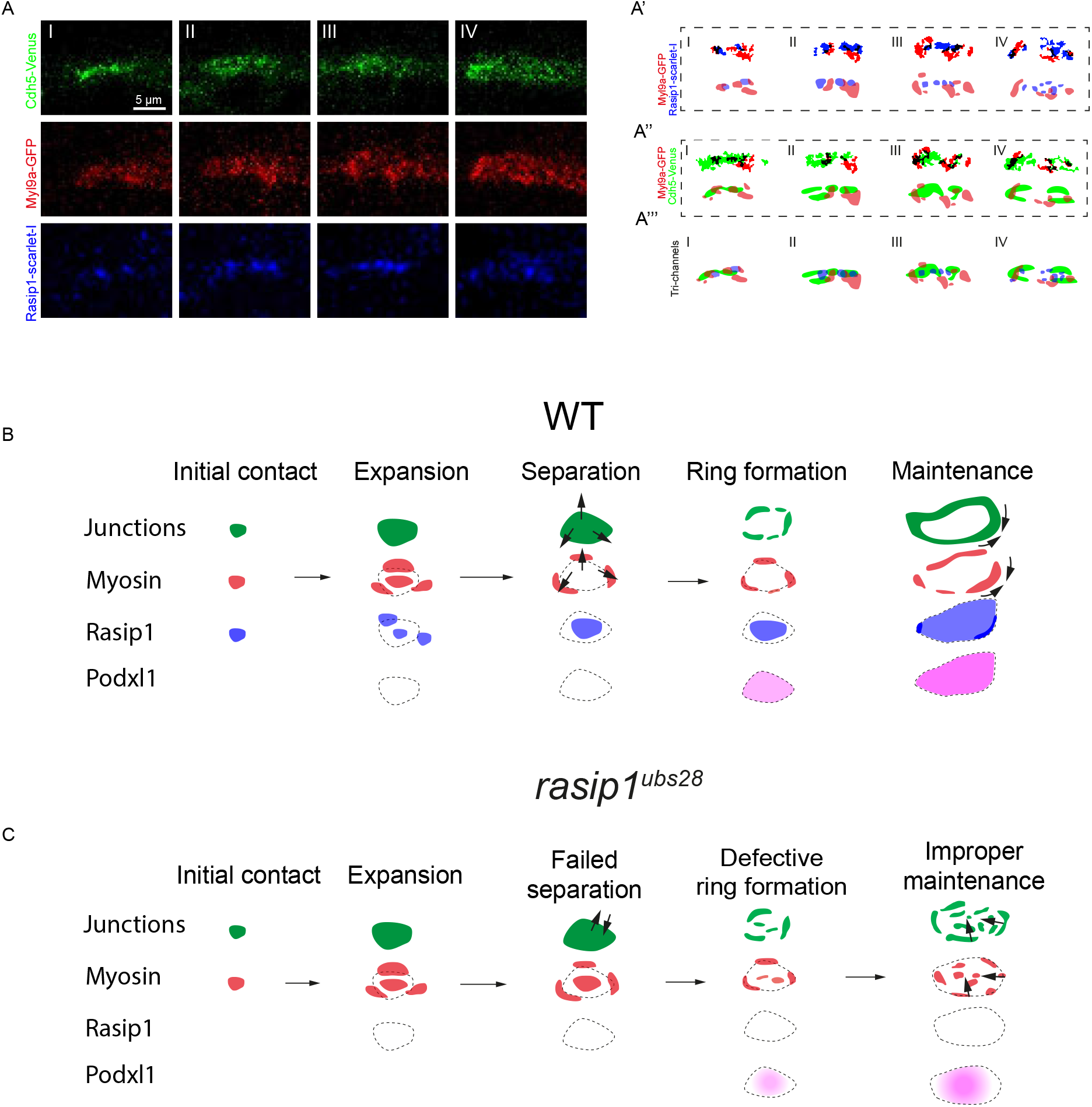
De-mixing of pre-apical patches and maintenance of apical compartments by delineating varying levels of actomyosin through Rasip1. (A)Time lapse of three-color live imaging showing the rearrangement of Cdh5, Myl9a and Rasip1 during *de novo* lumen formation. (A’) Automatic thresholding and diagrams showing the localization of Myl9a and Rasip1 clusters. (A’’) Automatic thresholding and diagrams showing the localization of Myl9a and Cdh5 clusters. (A’’’) Graphic diagrams showing the segregation between Cdh5, Myl9a and Rasip1 during *de novo* lumen formation. (B)Diagrams showing the molecule steps of *de novo* lumen formation. In wild-type embryos, both of Myosin and its inhibitor Rasip1 are recruited to the initial contact sites (initial contact). The initial contact sites expand into junctional patches (expansion), while Myosin and Rasip1 segregate and localize at the periphery and center respectively (Separation). Cdh5 relocates towards to the periphery with peripheralized contractility under the inhibition of Rasip1 in center (ring formation). After the junctional rings are formed, Rasip1 dynamically shuttles between the apical compartments and junctions tuning local contractility (maintenance). (C)In *rasip1* mutants, the pre-apical myosin fails to be inhibited, leading to unsustainable Cdh5 relocation towards the periphery (separation and ring formation). The ectopic apical myosin leads to radial contractility that de-stabilizes junctional proteins from boundaries (maintenance).

### Tuning contractility with the same pool of Rasip1 at distinct compartments

Canoe, a homologous protein of Rasip1 in Drosophila, is recruited to tricellular junctions of epithelial cells under high actomyosin contractility and tension (**Yu & Zallen, 2020**). The distinct responses of Canoe were proposed to respond to different levels of external forces, thus stabilizing cell-cell junctions or facilitating cell-cell rearrangements. During *de novo* lumen formation of the DLAV, Rasip1 relocates dynamically between the apical compartments and the junctions. Like Canoe, Rasip1 dynamically responds to high contractility at distinct cellular compartments by its varying localizations. The shuttling of Rasip1 between the apical compartments and junctions might convey another layer of dynamic regulation at compartment level. The dominant apical localization and rapid responsiveness at junctions of Rasip1 define and orchestrate different levels of actomyosin contractility in distinct compartments in a robust manner, utilizing the same pool of Rasip1. This occurs without the necessity of balancing two regulators at separate compartments. However, how Rasip1 and Canoe sense the high tension/contractility remains largely unknow. Further studies are required to explore the dynamic recruitment of Rasip1.

## Acknowledgements

We thank Kumuthini Kulendra for fish care and the Imaging Core Facility of the Biozentrum (University of Basel) for microscopy support. This work has been supported by the Kantons Basel-Stadt and Basel-Land and by grants from the Swiss National Science Foundation (310030_200701 and 310030B_176400) to M.A. We thank Hiroyuki Nakajima (NCVC, Osaka, Japan) for providing *Tg(fli1a: GFP-Podxl1)*^*ncv530Tg*^. We thank Anne Schmid (Institute Pasteur, Paris, France) for providing *Tg(kdrl: Myl9a-GFP)*^*ip5Tg*^. We thank the Michel Bagnat (Duke University, USA) for providing *TgKI(tjp1a-tdTomato)*^*pd1224*^.

## Author contributions

J.Y., H.-G.B. and M.A. designed the experiments. J.Y. performed the experiments and analyzed the data. J.Y. and M.L. analyzed the *rasip1* mutant phenotype. N.S., K.G. and J.Y. generated the *Tg(cdh5: cdh5-mClavGR2)*^*ubs58*^. J.Y., L.M. and M.P.K generated the *fli1a: RhoA-BcLOV4-mCherry* construct. L.-K.P. generated *Tg(fli:gal4; UAS: Myl9b-mCherry)*^*rk32*^. J.Y., C.W. and L.M. analyzed Myl9a distribution. J.Y., H.-G.B. and M.A. wrote the manuscript.

## References

Barry, D. M., Koo, Y., Norden, P. R., Wylie, L. A., Xu, K., Wichaidit, C., Azizoglu, D. B., Zheng, Y., Cobb, M. H., Davis, G. E., & Cleaver, O. (2016). Rasip1-Mediated Rho GTPase Signaling Regulates Blood Vessel Tubulogenesis via Nonmuscle Myosin II. Circulation Research, 119(7), 810–826. 10.1161/CIRCRESAHA.116.309094

Beckers, C. M. L., van Hinsbergh, V. W. M., & van Nieuw Amerongen, G. P. (2010). Driving Rho GTPase activity in endothelial cells regulates barrier integrity. Thrombosis and Haemostasis, 103(01), 40–55.

Berlew, E. E., Kuznetsov, I. A., Yamada, K., Bugaj, L. J., Boerckel, J. D., & Chow, B. Y. (2021). Single-Component Optogenetic Tools for Inducible RhoA GTPase Signaling. Advanced Biology, 5(9), 2100810. 10.1002/adbi.202100810

Betz, C., Lenard, A., Belting, H.-G., & Affolter, M. (2016). Cell behaviors and dynamics during angiogenesis. Development, 143(13), 2249–2260. 10.1242/dev.135616

Blum, Y., Belting, H.-G., Ellertsdo|r, E., Herwig, L., Lüders, F., & Affolter, M. (2008). Complex cell rearrangements during intersegmental vessel sprouting and vessel fusion in the zebrafish embryo. Developmental Biology, 316(2), 312–322. 10.1016/j.ydbio.2008.01.038

Daia, A., Bryant, D. M., & Mostov, K. E. (2011). Molecular Regulation of Lumen Morphogenesis. Current Biology, 21(3), R126–R136. 10.1016/j.cub.2010.12.003

Davis, G. E., Stratman, A. N., Sacharidou, A., & Koh, W. (2011). Chapter Three - Molecular Basis for Endothelial Lumen Formation and Tubulogenesis During Vasculogenesis and Angiogenic Sprouting. In K. W. Jeon (Ed.), International Review of Cell and Molecular Biology (Vol. 288, pp. 101–165). Academic Press. 10.1016/B978-0-12-386041-5.00003-0

Dorland, Y. L., & Huveneers, S. (2017). Cell–cell junctional mechanotransduction in endothelial remodeling. Cellular and Molecular Life Sciences, 74(2), 279–292. 10.1007/s00018-016-2325-8

Eelen, G., Treps, L., Li, X., & Carmeliet, P. (2020). Basic and Therapeutic Aspects of Angiogenesis Updated. Circulation Research, 127(2), 310–329. 10.1161/CIRCRESAHA.120.316851

Herwig, L., Blum, Y., Krudewig, A., Ellertsdo|r, E., Lenard, A., Belting, H. G., & Affolter, M. (2011). Distinct cellular mechanisms of blood vessel fusion in the zebrafish embryo. Current Biology, 21(22), 1942–1948. 10.1016/j.cub.2011.10.016

Honigmann, A., Sadeghi, S., Keller, J., Hell, S. W., Eggeling, C., & Vink, R. (2014). A lipid bound actin meshwork organizes liquid phase separation in model membranes. ELife, 3, e01671. 10.7554/eLife.01671

Iruela-Arispe, M. L., & Davis, G. E. (2009). Cellular and Molecular Mechanisms of Vascular Lumen Formation. Developmental Cell, 16(2), 222–231. 10.1016/j.devcel.2009.01.013

Koo, Y., Barry, D. M., Xu, K., Tanigaki, K., Davis, G. E., Mineo, C., & Cleaver, O. (2016). Rasip1 is essential to blood vessel stability and angiogenic blood vessel growth. Angiogenesis, 19(2), 173–190. 10.1007/s10456-016-9498-5

Krapf, D. (2018). Compartmentalization of the plasma membrane. Current Opinion in Cell Biology, 53, 15–21. 10.1016/j.ceb.2018.04.002

Lagendijk, A. K., Gomez, G. A., Baek, S., Hesselson, D., Hughes, W. E., Paterson, S., Conway, D. E., Belting, H. G., Affolter, M., Smith, K. A., Schwartz, M. A., Yap, A. S., & Hogan, B. M. (2017). Live imaging molecular changes in junctional tension upon VE-cadherin in zebrafish. Nature Communications, 8(1), 1402. 10.1038/s41467-017-01325-6

Lancino, M., Majello, S., Herbert, S., De Chaumont, F., Tinevez, J.-Y., Olivo-Marin, J.-C., Herbomel, P., & Schmidt, A. (2018). Anisotropic organization of circumferential actomyosin characterizes hematopoietic stem cells emergence in the zebrafish. ELife, 7, e37355. 10.7554/eLife.37355

Lawson, N. D., & Weinstein, B. M. (2002). In Vivo Imaging of Embryonic Vascular Development Using Transgenic Zebrafish. Developmental Biology, 248(2), 307–318. 10.1006/dbio.2002.0711

Lee, M., Betz, C., Yin, J., Paatero, I., Schellinx, N., Carte, A. N., Wilson, C. W., Ye, W., Affolter, M., & Belting, H. G. (2021). Control of dynamic cell behaviors during angiogenesis and anastomosis by Rasip1. Development (Cambridge), 148(10). 10.1242/DEV.197509/VIDEO-14

Levic, D. S., Yamaguchi, N., Wang, S., Knaut, H., & Bagnat, M. (2021). Knock-in tagging in zebrafish facilitated by insertion into non-coding regions. Development, 148(19), dev199994. 10.1242/dev.199994

Post, A., Pannekoek, W. J., Ponsioen, B., Vliem, M. J., & Bos, J. L. (2015). Rap1 Spatially Controls ArhGAP29 To Inhibit Rho Signaling during Endothelial Barrier Regulation. Molecular and Cellular Biology, 35(14), 2495–2502. 10.1128/MCB.01453-14

Post, A., Pannekoek, W.-J., Ross, S. H., Verlaan, I., Brouwer, P. M., & Bos, J. L. (2013). Rasip1 mediates Rap1 regulation of Rho in endothelial barrier function through ArhGAP29. Proceedings of the National Academy of Sciences, 110(28), 11427–11432. 10.1073/pnas.1306595110

Robbins, R. M., & Beitel, G. J. (2010). Vascular Lumen Formation: Negativity Will Tear Us Apart. Current Biology, 20(22), R973–R975. 10.1016/j.cub.2010.10.032

Schuermann, A., Helker, C. S. M., & Herzog, W. (2014). Angiogenesis in zebrafish. Seminars in Cell & Developmental Biology, 31, 106–114. 10.1016/j.semcdb.2014.04.037

Sigurbjörnsdó|r, S., Mathew, R., & Leptin, M. (2014). Molecular mechanisms of de novo lumen formation. Nature Reviews Molecular Cell Biology, 15(10), 665–676. 10.1038/nrm3871

Strilic, B., Eglinger, J., Krieg, M., Zeeb, M., Axnick, J., Babál, P., Müller, D. J., & Lammert, E. (2010). Electrostatic Cell-Surface Repulsion Initiates Lumen Formation in Developing Blood Vessels. Current Biology, 20(22), 2003–2009. 10.1016/j.cub.2010.09.061

Strilic, B., Kučera, T., Eglinger, J., Hughes, M. R., McNagny, K. M., Tsukita, S., Dejana, E., Ferrara, N., & Lammert, E. (2009). The Molecular Basis of Vascular Lumen Formation in the Developing Mouse Aorta. Developmental Cell, 17(4), 505–515. 10.1016/j.devcel.2009.08.011

Thielicke, W., & Sonntag, R. (2021). Particle Image Velocimetry for MATLAB: Accuracy and enhanced algorithms in PIVlab. Journal of Open Research Software. 10.5334/jors.334

Vogel, S. K., Greiss, F., Khmelinskaia, A., & Schwille, P. (2017). Control of lipid domain organization by a biomimetic contractile actomyosin cortex. ELife, 6, e24350. 10.7554/eLife.24350

Wilson, C. W., Parker, L. H., Hall, C. J., Smyczek, T., Mak, J., Crow, A., Posthuma, G., De Mazière, A., Sagolla, M., Chalouni, C., Vitorino, P., Roose-Girma, M., Warming, S., Klumperman, J., Crosier, P. S., & Ye, W. (2013). Rasip1 regulates vertebrate vascular endothelial junction stability through Epac1-Rap1 signaling. Blood, 122(22), 3678–3690. 10.1182/blood-2013-02-483156

Xu, K., Chong, D. C., Rankin, S. A., Zorn, A. M., & Cleaver, O. (2009). Rasip1 is required for endothelial cell motility, angiogenesis and vessel formation. Developmental Biology, 329(2), 269–279. 10.1016/j.ydbio.2009.02.033

Xu, K., & Cleaver, O. (2011). Tubulogenesis during blood vessel formation. Seminars in Cell & Developmental Biology, 22(9), 993–1004. 10.1016/j.semcdb.2011.05.001

Xu, K., Sacharidou, A., Fu, S., Chong, D. C., Skaug, B., Chen, Z. J., Davis, G. E., & Cleaver, O. (2011). Blood Vessel Tubulogenesis Requires Rasip1 Regulation of GTPase Signaling. Developmental Cell, 20(4), 526–539. 10.1016/j.devcel.2011.02.010

Yu, H. H., & Zallen, J. A. (2020). Abl and Canoe/Afadin mediate mechanotransduction at tricellular junctions. Science, 370(6520), eaba5528. 10.1126/science.aba5528

