## Supplementary for "Initiation of Lumen Formation from Junctions via Differential Actomyosin Contractility Regulated by Dynamic Recruitment of Rasip1"

**Supplementary Experimental Procedures**
**Supplementary Note**
**Supplementary References**
**Figure S1**, related to Figure 1
**Figure S2**, related to Figure 2
**Figure S3**, related to Figure 3
**Figure S4**, related to Figure 4
**Figure S5**, related to Figure 5
**Figure S6**, related to Figure 6
**Figure S7**, related to Figure 7
**Figure S8**, related to Supplementary Experimental Procedures
**Video S1,** related to Figure 1A
**Video S2,** related to Figure 1B
**Video S3,** related to Figure 1C
**Video S4,** related to Figure 2I
**Video S5,** related to Figure 3E’
**Video S6,** related to Figure 3G’
**Video S7,** related to Figure 4A
**Video S8,** related to Figure 4B
**Video S9,** related to Figure 4C
**Video S10,** related to Figure S4B
**Video S11,** related to Figure 5A
**Video S12,** related to Figure 5C
**Video S13,** related to Figure 5G
**Video S14,** related to Figure 5H
**Video S15,** related to Figure 6D and 6E
**Video S16,** related to Figure 6F
**Video S17,** related to Figure 6G
**Video S18,** related to Figure 6I
**Video S19,** related to Figure 6J
**Video S20,** related to Figure 7H

### Supplementary Experimental Procedures

**Fish maintenance and stocks.** Zebrafish (*Danio rerio*) were maintained according to FELASA guidelines (Aleström et al., 2019). All experiments were performed in accordance with federal guidelines and were approved by the Kantonales Veterinäramt of Kanton Basel-Stadt (1027H, 1014HE2, 1014G). New zebrafish lines developed in this study: *Tg(cdh5: cdh5-mClavGR2)*$^{ubs58}$, *Tg(fli:gal4; UAS: Myl9b-mCherry)*$^{rk32}$, *Tg(fli1a: Rasip1-scarlet-I)*$^{ubs59}$. Previously established zebrafish lines: *Tg(cdh5: cdh5-TFP-TENS-Venus)*$^{uq11bh}$ (Lagendijk et al., 2017), *TgKI(tjp1a-tdTomato)*$^{pd1224}$ (Levic et al., 2021), *Tg(fli1a: GFP-Podxl1)* $^{ncv530Tg}$, *Tg(kdrl: Myl9a-GFP)*$^{ip5Tg}$ (Lancino et al., 2018), *Tg(fli1ep:gal4ff)*$^{ubs3}$ (Zygmunt et al., 2011), *Tg(UAS:mRuby2-UCHD)*$^{ubs20}$ (Paatero et al., 2018), *rasip1*$^{ubs28}$ (Lee et al., 2021) Transient expressions in zebrafish embryos: *fli1a: GFP-Rasip1*, *fli1a: RhoA-BcLOV4-mCherry*.

**Molecular biology and transgenesis.** All plasmids were generated through Gibson assembly, Gateway or restriction cloning unless otherwise stated. All constructs were cloned into tol2 vectors before injections into zebrafish embryos unless otherwise stated. All details for plasmid cloning and transgenesis were in Supplementary Note 1.

**Live imaging, photo-activation and optogenetics.** Zebrafish embryos between 30 hpf to 32 hpf were dechorionated and anaesthetized with 0.16 mg ml$^{-1}$ (1×) tricaine methanesulfonate (Sigma). Embryos were mounted into microwell dishes within 0.75% low-melting-point agarose (ROTI) and were covered with E3 buffer with 1× tricaine. Live imaging, photo-conversion and opto-activation were performed with Leica SP5 confocal microscope with ×40 water immersion objective. Photo-conversions were applied at manually selected region of interest (ROI) with 405nm laser for 10-20s before subsequent live imaging. Conversion at ROI was monitored with simultaneous imaging at 488nm to guarantee complete conversion. Optogenetic activations were carried out at manually selected ROI with 450nm laser continuously throughout live imaging. Sequential imaging of Venus and GFP was described in Figure S8.

**Image analysis and Data analysis.** Images were analysed using the FiJi and MATLAB with home-made codes. Z-stacks were flattened by maximum or sum slices intensity projections in Fiji software. Correction of drift of images, PIV, quantification of relative signal levels at distinct cellular compartments were carried out with MATLAB with home-made codes. Image panels were made with Open Microscopy Environment (OMERA). MATLBA was used for all statistical analysis (*t* test). No statistical methods were used to predetermine sample sizes.

**Immunofluorescence**: For immunofluorescence and imaging of zebrafish embryos, dechorionated zebrafish embryos were fixed in 2% paraformaldehyde and 0.1% Tween 20 in PBS (PBST) overnight at 4 °C unless otherwise stated. For immunofluorescence involved Rasip1, dechorionated embryos were fixed in 2% TCA in PBST for 15min at room temperature. After fixation, embryos were washed four times with PBST and permeabilized with 0.5% Triton-X-100 in PBS at room temperature (RT) for 30 minutes and were blocked with 2% BSA and 5% goat serum in PSBT overnight with continuous shaking. The following antibodies were used: rabbit anti-zf-Podxl (1:200) (Herwig et al., 2011), rabbit anti-zf-Cdh5 (1:200) (Blum et al., 2008), guinea pigs anti-zf-Cdh5 (1:200), rabbit anti-Rasip1 (1:500) (Lee et al., 2021), mouse anti-human ZO-1 (Invitrogen, # 33-9100,1:500). Embryos were incubated with primary and secondary antibodies at 4 °C overnight with continuous shaking respectively and washed 6 times in between. After immunostaining, embryos were mounted within 0.75% low-melting-point agarose and images were taken with Leica SP5 with 40x water immersion objective.

**Chemical treatments.** Embryos were incubated in E3 medium containing 75 µM Y-27632 (MCE, HY-10583) or 1% DMSO as controls. The same concentrations of chemicals were applied to the low-melting-point agarose mounting medium and the E3 medium on top of it before imaging.

#### Supplementary Note 1: Molecular biology and transgenesis

##### 1.1 Cloning and transgenesis of *Tg(cdh5: cdh5-mClavGR2)*$^{ubs58}$

The sequence encoding for *mClavGR2* was introduced by BAC recombineering technology into the exon 13 of cdh5 between the p120-catenin and β-catenin binding sites of the translated Cdh5 protein. This position has been demonstrated harmless to the proper function of Cdh5 (Lagendijk et al., 2017). Defective λ prophage system was used for BAC recombineering (Sharan et al., 2009). The BAC CH73-357K2 containing the gene of *Cdh5* was transformed into the SW102 E. coli. BAC containing SW102 cells were then inoculated and transformed with the *galK* gene flanked by homology arms. One surviving colony was subsequently inoculated and transformed with the insert containing *mClavGR2*, flanked by the same homology arms as before. Transformed cells were selected on DOG containing minimal plates (negative selection for galK expression). Surviving colonies were analysed for desired modification. The selected *cdh5-mClav* containing BAC was subsequently tested by XbaI digest, further PCR analyses and sequencing.

Galk_mClav_fo: TGGCCATGATGATCGAGGTGAAGAAGGACGAGGCAGATCGTGATCGAGATCCTGTTGACAATTAATCATC GGCA

Galk_mClav_re :GATTCGGTTCCCTCATAGCCATAGATGTGTAATGTGTCATAGGGAATGCCTCAGCACTGTCCTGCTCCTT

Homo_mClav_fo:TGGCCATGATGATCGAGGTGAAGAAGGACGAGGCAGATCGTGATCGAGATGCCGGCGTGAGCAAGGG CGAGGAG

Homo_mClav_re:GATTCGGTTCCCTCATAGCCATAGATGTGTAATGTGTCATAGGGAATGCCGCCGGCCTTGTACAGCTCGT CCATGCC

##### 1.2 Cloning and transgenesis of *Tg(fli1a:Rasip1-scarlet-I)*$^{ubs59}$

The cDNA of *Rasip1* was cloned from zebrafish cDNA library into pME vector through the BP reaction of gateway cloning.

rasip1-BP-F: GGGGACAAGTTTGTACAAAAAAGCAGGCTCTCGAGatggaggagtctggcagtc

rasip1-BP-R: GGGGACCACTTTGTACAAGAAAGCTGGGTttacagtctcgtccgct

The restrictive sites of NotI and MfeI were introduced into *pME-Rasip1* through PCR at the 3' end of *Rasip1* cDNA.

pME-Rasip1-3R-NotI-F: AATCATGCGCGGCCGCTAAAACCCAGCTTTCTTGTACAAAGTTGGCATTATAAG

pME-Rasip1-3R-MfeI-R: ATACATACCAATTGGCCTCCGGATCCGCCCAGTCTCGTCCGCTGATTAG

The cDNA of *scarlet-I* was synthesized from Genewiz with codon optimized for zebrafish. Restriction-ligation was used for insertion of Scarlet-I to the c-terminal of Rasip1 with linker: GGSGGQL.

Scarlet-I-F: ATACGTGGCAATTGATGGTGAGCAAGGGCGAG

Scarlet-I-R: TTATCCAAGCGGCCGCCGCTTCCTCCTCCGCTTCCTCCcttgtacagctcgtccat

The constructed *pME-Rasip1-scarlet-I* was fully sequenced. The *pME-Rasip1-scarlet-I* was cloned into PDestTol2CG2 through the LR reaction of gateway cloning with a 2kb *fli1a* promoter and 3' poly A tail.

**1.3 Cloning of *fli1a:GFP-Rasip1***

*GFP* was inserted to the 5' end of *Rasip1* into the *pME-Rasip1* vector with linker: GGSGGGSGG through Gibson assembly.

GFP-Gib-F:GTACAAAAAAGCAGGCTCATGGTGAGCAAGGGCGAG

GFP-Gib-R: ctgccagactcctccatCCCTCCGCTTCCTCCTCCGCTTCCTCCCTTGTACAGCTCGTCCATGC

pME-rasip1-Gib-F: Gatggaggagtctggcagtcc

pME-rasip1-Gib-R; GCCTGCTTTTTTGTACAAAGTTGGCAT

The constructed *pME-GFP-Rasip1* was fully sequenced. The *pME-GFP-Rasip1* was cloned into PDestTol2CG2 through the LR reaction of gateway cloning with a 2kb fli1a promoter and 3' poly A tail.

**1.4 Cloning of *fli1a: RhoA-BcLOV4-mCherry***

the light-oxygen-voltage (LOV) flavoprotein BcLOV4, derived from Botrytis cinerea, swiftly relocates to the plasma membrane through an electrostatic interaction with lipids on the inner leaflet, a process regulated by blue light (Glantz et al., 2018, 2019). It was used as a single-component optogenetic tool for inducible RhoA GTPase Signaling in cell culture (Berlew et al., 2021). We ordered the opto-RhoA-mCherry_pcDNA3.1 construct from addgene (#164472) .

The Tol2 vector PDestTol2CG2 carried a 2Kb fli1a promoter was amplified with PCR to gain the restriction sites EcoRI and NheI. *RhoA-BcLOV4-mCherry* was cloned into Tol2 destination vector PDestTol2CG2 by adding restriction sites MfeI and XbaI through PCR.

tol2-NheI-F: GATTTTCTGCTAGCggtaccatcgatgatgatccag

tol2-fli-EcoRI-R:TTAGTACTGAATTCAATTCCACCGCGTCTGAATTAATTCCAGCC

opto-RhoA-mfeI-F: GTCAGTCACAATTGatgtcagctgccatccgg

opto-RhoA-xbaI-R: ACTGACTGTCTAGAttacttgtacagctcgtccatgc

#### Supplementary Reference:


Aleström, P., D'Angelo, L., Midtlyng, P. J., Schorderet, D. F., Schulte-Merker, S., Sohm, F., & Warner, S. (2019). Zebrafish: Housing and husbandry recommendations. *Laboratory Animals*, *54*(3), 213–224. https://doi.org/10.1177/0023677219869037

Berlew, E. E., Kuznetsov, I. A., Yamada, K., Bugaj, L. J., Boerckel, J. D., & Chow, B. Y. (2021). Single-Component Optogenetic Tools for Inducible RhoA GTPase Signaling. *Advanced Biology*, *5*(9), 2100810. https://doi.org/https://doi.org/10.1002/adbi.202100810

Blum, Y., Belting, H.-G., Ellertsdottir, E., Herwig, L., Lüders, F., & Affolter, M. (2008). Complex cell rearrangements during intersegmental vessel sprouting and vessel fusion in the zebrafish embryo. *Developmental Biology*, *316*(2), 312–322. https://doi.org/https://doi.org/10.1016/j.ydbio.2008.01.038

Glantz, S. T., Berlew, E. E., & Chow, B. Y. (2019). Chapter Eleven - Synthetic cell-like membrane interfaces for probing dynamic protein-lipid interactions. In A. K. Shukla (Ed.), *Methods in Enzymology* (Vol. 622, pp. 249–270). Academic Press. https://doi.org/https://doi.org/10.1016/bs.mie.2019.02.015

Glantz, S. T., Berlew, E. E., Jaber, Z., Schuster, B. S., Gardner, K. H., & Chow, B. Y. (2018). Directly light-regulated binding of RGS-LOV photoreceptors to anionic membrane phospholipids. *Proceedings of the National Academy of Sciences*, *115*(33), E7720–E7727. https://doi.org/10.1073/pnas.1802832115

Herwig, L., Blum, Y., Krudewig, A., Ellertsdottir, E., Lenard, A., Belting, H. G., & Affolter, M. (2011). Distinct cellular mechanisms of blood vessel fusion in the zebrafish embryo. *Current Biology*, *21*(22), 1942–1948. https://doi.org/10.1016/j.cub.2011.10.016

Hogan, B. M., Bussmann, J., Wolburg, H., & Schulte-Merker, S. (2008). ccm1 cell autonomously regulates endothelial cellular morphogenesis and vascular tubulogenesis in zebrafish. *Human Molecular Genetics*, *17*(16), 2424–2432. https://doi.org/10.1093/hmg/ddn142

Lagendijk, A. K., Gomez, G. A., Baek, S., Hesselson, D., Hughes, W. E., Paterson, S., Conway, D. E., Belting, H. G., Affolter, M., Smith, K. A., Schwartz, M. A., Yap, A. S., & Hogan, B. M. (2017). Live imaging molecular changes in junctional tension upon VE-cadherin in zebrafish. *Nature Communications*, *8*(1), 1402. https://doi.org/10.1038/s41467-017-01325-6

Lancino, M., Majello, S., Herbert, S., De Chaumont, F., Tinevez, J.-Y., Olivo-Marin, J.-C., Herbomel, P., & Schmidt, A. (2018). Anisotropic organization of circumferential actomyosin characterizes hematopoietic stem cells emergence in the zebrafish. *ELife*, *7*, e37355. https://doi.org/10.7554/eLife.37355

Lee, M., Betz, C., Yin, J., Paatero, I., Schellinx, N., Carte, A. N., Wilson, C. W., Ye, W., Affolter, M., & Belting, H. G. (2021). Control of dynamic cell behaviors during angiogenesis and anastomosis by Rasip1. *Development (Cambridge)*, *148*(10). https://doi.org/10.1242/DEV.197509/VIDEO-14

Levic, D. S., Yamaguchi, N., Wang, S., Knaut, H., & Bagnat, M. (2021). Knock-in tagging in zebrafish facilitated by insertion into non-coding regions. *Development*, *148*(19), dev199994. https://doi.org/10.1242/dev.199994

Mably, J. D., Burns, C. G., Chen, J.-N., Fishman, M. C., & Mohideen, M.-A. P. K. (2003). heart of glass Regulates the Concentric Growth of the Heart in Zebrafish. *Current Biology*, *13*(24), 2138–2147. https://doi.org/https://doi.org/10.1016/j.cub.2003.11.055

Paatero, I., Sauteur, L., Lee, M., Lagendijk, A. K., Heutschi, D., Wiesner, C., Guzmán, C., Bieli, D., Hogan, B. M., Affolter, M., & Belting, H. G. (2018). Junction-based lamellipodia drive endothelial cell rearrangements in vivo via a VE-cadherin-F-actin based oscillatory cell-cell interaction. *Nature Communications*, *9*(1), 3545. https://doi.org/10.1038/s41467-018-05851-9

Sauteur, L., Krudewig, A., Herwig, L., Ehrenfeuchter, N., Lenard, A., Affolter, M., & Belting, H.-G. (2014). Cdh5/VE-cadherin Promotes Endothelial Cell Interface Elongation via Cortical Actin Polymerization during Angiogenic Sprouting. *Cell Reports*, *9*(2), 504–513. https://doi.org/https://doi.org/10.1016/j.celrep.2014.09.024

Sharan, S. K., Thomason, L. C., Kuznetsov, S. G., & Court, D. L. (2009). Recombineering: a homologous recombination-based method of genetic engineering. *Nature Protocols*, *4*(2), 206–223. https://doi.org/10.1038/nprot.2008.227

Zygmunt, T., Gay, C. M., Blondelle, J., Singh, M. K., Flaherty, K. M., Means, P. C., Herwig, L., Krudewig, A., Belting, H.-G., Affolter, M., Epstein, J. A., & Torres-Vázquez, J. (2011). Semaphorin-PlexinD1 Signaling

Limits Angiogenic Potential via the VEGF Decoy Receptor sFlt1. *Developmental Cell*, *21*(2), 301–314. https://doi.org/https://doi.org/10.1016/j.devcel.2011.06.033

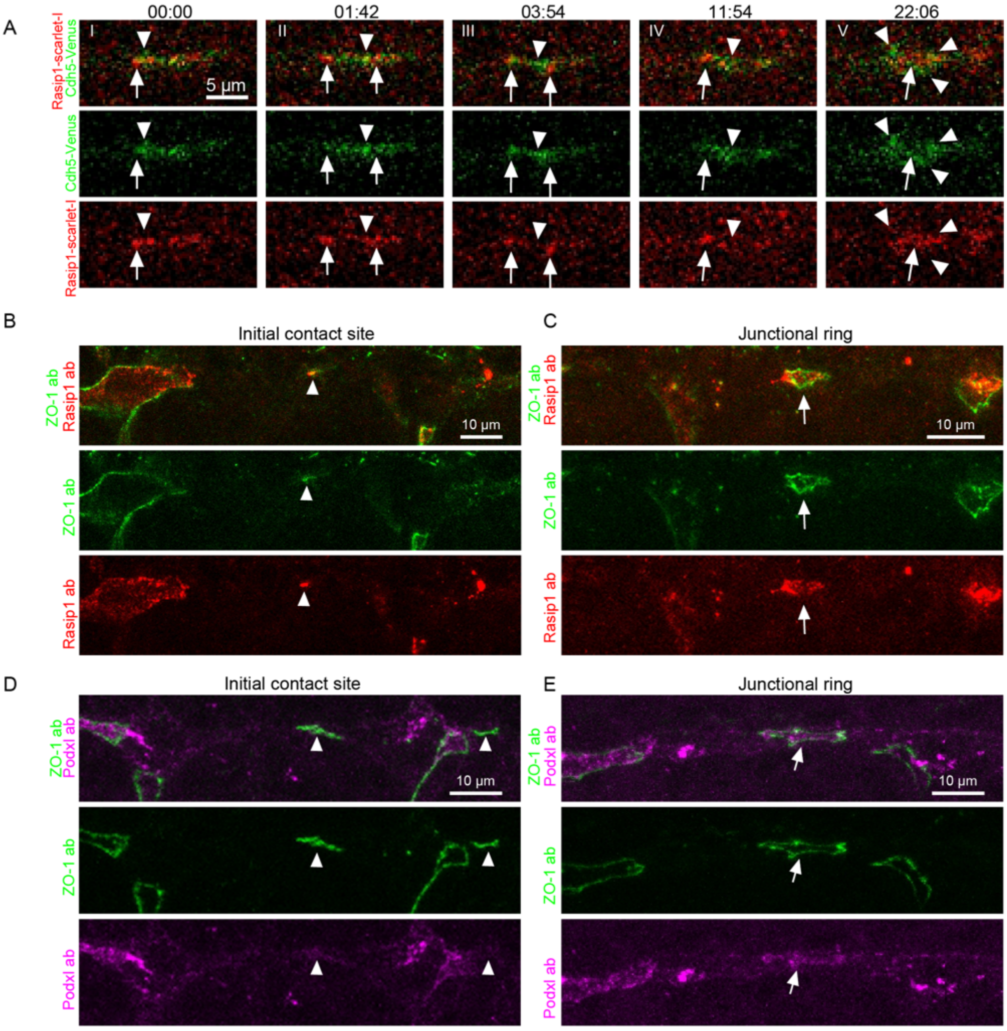


**Figure S1. Recruitment of Rasip1 and Podxl1 during the patch-to-ring transition, related to Figure 1**

(A) Time lapses showing the recruitment of Rasip1 to the junctional patches and nascent junctional rings. Rasip1 translocated to the periphery of the junctional patches. White arrows label Cdh5 whilst white arrowheads label Rasip1.

(B and C) Antibody staining of ZO-1 and Rasip1 at junctional patches (B) or nascent junctional rings (C).

(D and E) Antibody staining of ZO-1 and Podxl at junctional patches (D) or nascent junctional rings (E). White arrowheads label the junctional patches whilst white arrows label the nascent junctional rings formed by anastomosis.

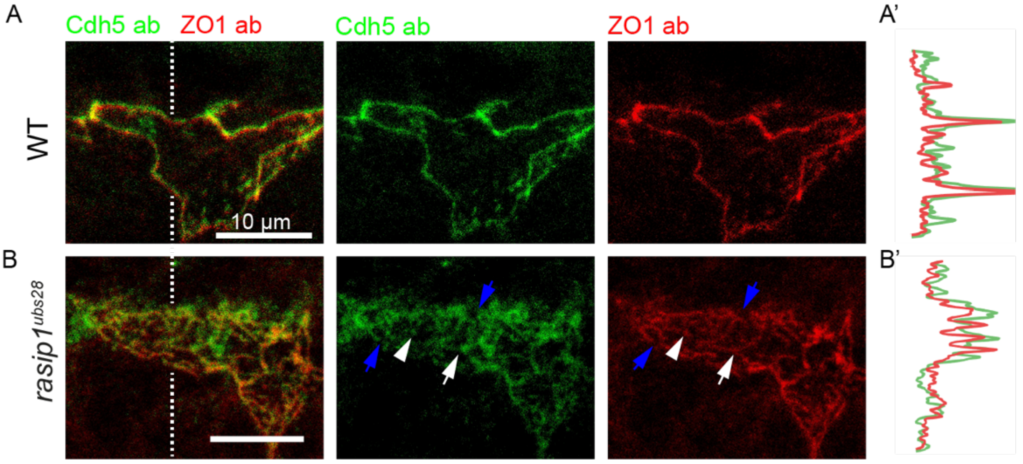


**Figure S2. Junctional materials are mis-localized in *rasip1$^{ubs28}$* mutants from proper junctions, related to Figure 2**

(A and B) Antibody staining of Cdh5 and ZO-1 in WT embryos and *rasip1$^{ubs28}$* mutants. Blue arrows label discontinuous junctions. White arrowheads and arrows label junctional clusters or linear junctional structures respectively. (A' and B') Intensities of Cdh5 antibody and ZO1 antibody along the dashed lines in (A) and (B).

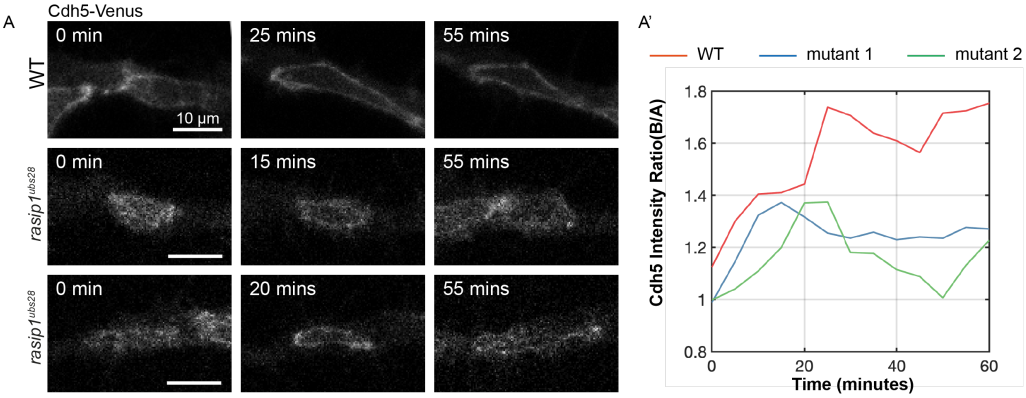


**Figure S3. Cdh5 clusters detach from junctions and move towards the apical compartments in *rasip1^ubs28^* mutants, related to Figure 3**

(A) Time lapses of Cdh5-Venus in WT embryos and *rasip1*$^{ubs28}$ mutants. The junctional patches transited into rings transiently but collapsed soon in *rasip1*$^{ubs28}$ mutants. (A') Temporal profiles of the Cdh5 boundary-to-apical ratio in WT embryos and *rasip1*$^{ubs28}$ mutants in (A). Mutant 1: middle panel of (A); Mutant 2: bottom panel of (A).

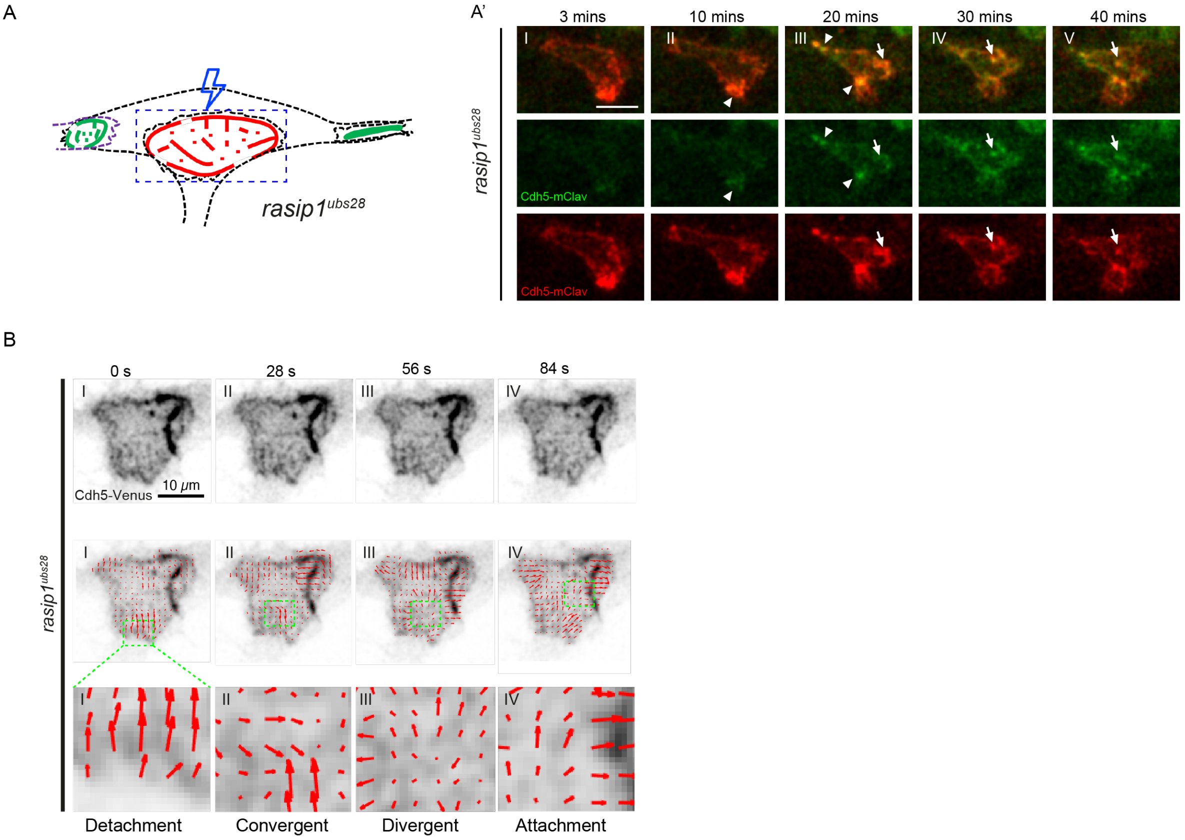


**Figure S4. Dynamics of Cdh5 measured by photo-conversion and PIV, related to Figure 4**

(A and A') Diagram and time-lapse of photo-conversion on the junctional ring between tip and stalk cells in *rasip1*$^{ubs28}$ mutants. Photo-conversion was performed before the degeneration of the junctional ring into reticulated junctions. White arrowheads label newly incorporated green Cdh5-mClav at the boundaries. White arrows label the Cdh5 clusters moving towards the apical compartment.

(B)Time lapse of Cdh5-Venus in *rasip1*$^{ubs28}$ mutants and corresponding quiver plots generated by PIV showing different patterns of Cdh5 rearrangement at the apical compartments.

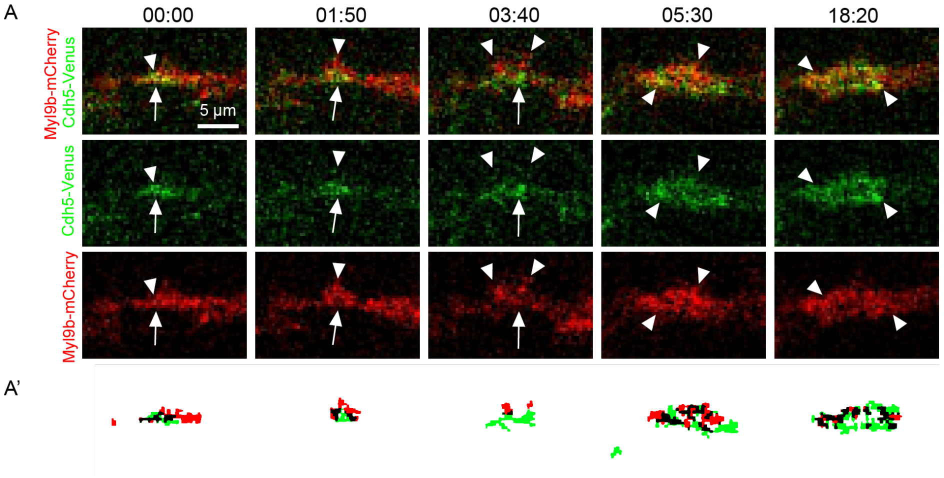


**Figure S5. Rearrangement of Myl9b and Cdh5 during the patch-to-ring transition or at junctions, related to Figure 5**

(A)Time lapses of Cdh5-Venus and Myl9b-mCherry showing the rearrangement of Myl9b through the patch-ring transition. White arrowheads label Myl9b clusters whilst white arrows label Cdh5 patches. (A’) Automatic thresholding of Myl9b-mCherry and Cdh5-Venus in (A).

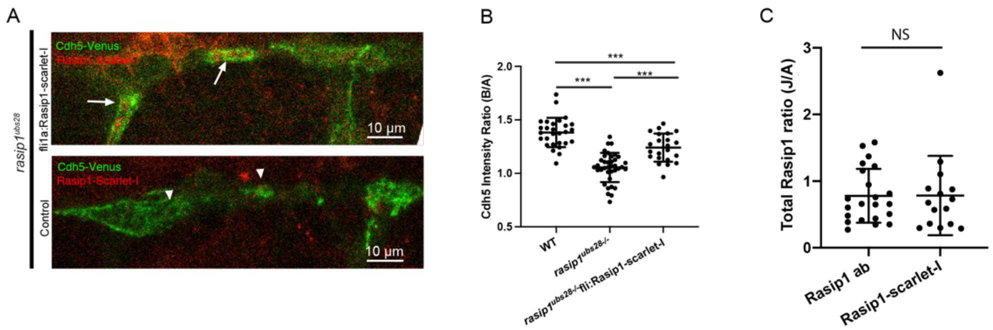


**Figure S6. Rasip1 reporters rescued the apical clearance defects and displayed a similar pattern of localization at junctions and within the apical compartments compared to antibody staining, related to Figure 6**

(A and B) Expression of Rasip1-scarlet-I under *fli1a* promoter rescued the apical clearance phenotype in *rasip1$^{ubs28}$* mutants (WT: n=28; *rasip1$^{ubs28}$* : n=37; *rasip1$^{ubs28}$* +Rasip1-scarlet-I: n=22). White arrows label the opened junctional rings with expression of Rasip1-scarlet-I. White arrowheads label the unopened or reticulated junctions in the absence of Rasip1-scarlet-I in *rasip1$^{ubs28}$* mutants.

(C) Quantification of the boundary-to-apical ratio of the total amount of Rasip1 in each compartment based on Rasip1 antibody or the Rasip1 live reporter.

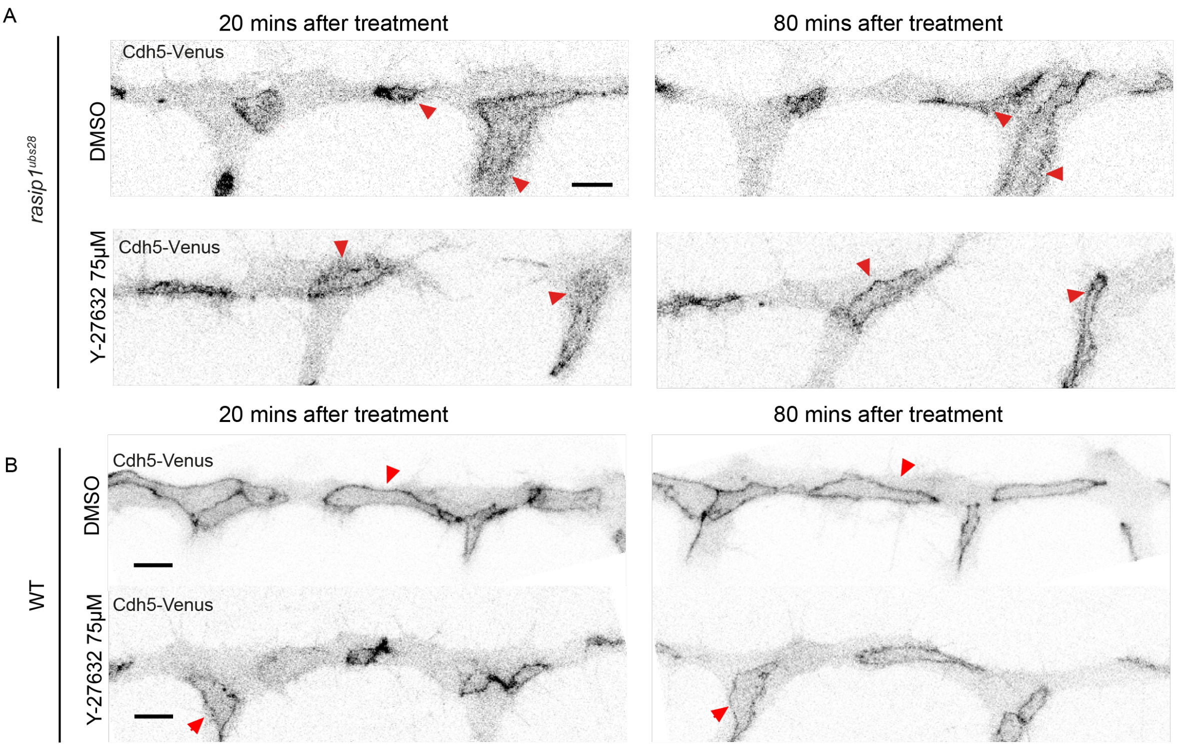


**Figure S7. Inhibition of ROCK alleviates the ectopic apical Cdh5 in *rasip1*$^{ubs28}$ mutants, related to Figure 7**

(A) Cdh5-Venus at 20 minutes and 80 minutes after 1% DMSO or Y-27632(75 µM) treatment in *rasip1*$^{ubs28}$ mutants. Red arrowheads exhibit the same junctions at 20 minutes and 80 minutes after drug treatment.

(B) Cdh5-Venus at 20 minutes and 80 minutes after 1% DMSO or Y-27632(75 µM) treatment in WT embryos. Red arrowheads exhibit the same junctions at 20 minutes and 80 minutes after drug treatment.

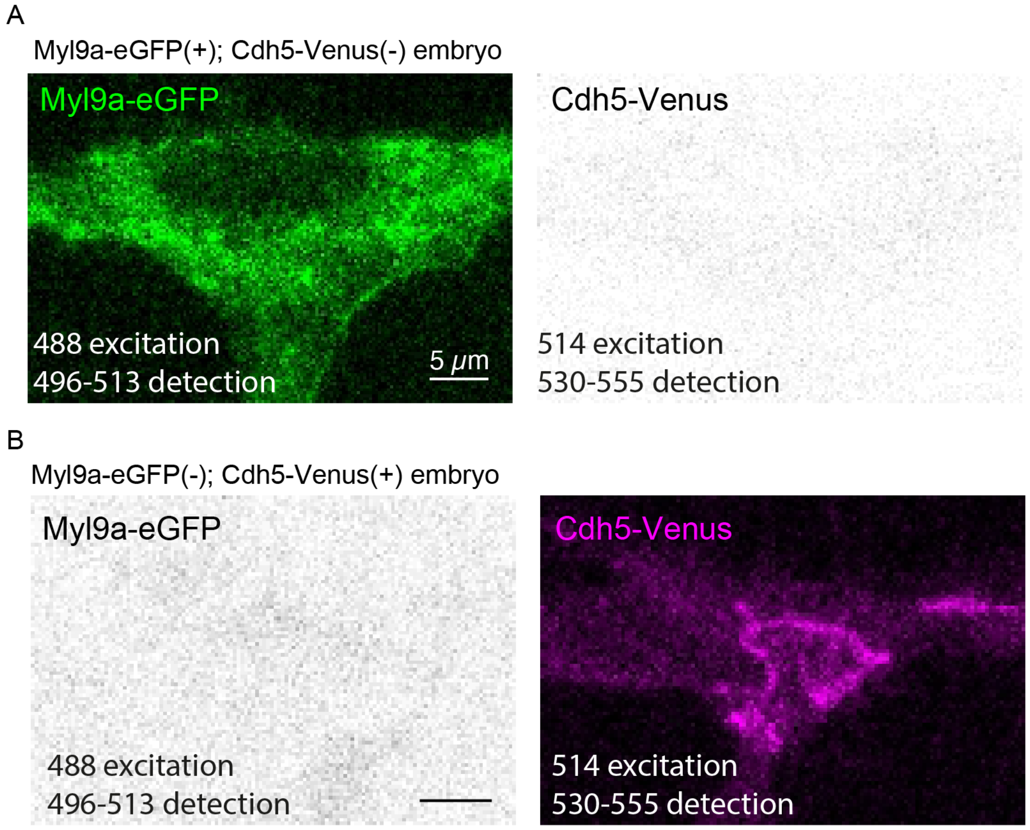


**Figure S8. Test of possible crosstalk during live imaging of Myl9a-eGFP and Cdh5-Venus, related to Supplementary Experimental Procedures**

(A) Myl9a-eGFP was excited by 488 nm laser and detected at the range between 496 nm and 513 nm in embryos that are only Myl9a-eGFP positive.

(B) Cdh5-Venus was excited by 514 nm laser and detected at the range between 530 nm and 555 nm in embryos that are only Cdh5-Venus positive.

**Supplementary Video legends**

**Video S1** (related to Figure 1A). Anastomosis and *de novo* lumen formation in zebrafish DLAV. Time-lapse series with expression of Cdh5-Venus (green) and GFP-Podxl1 (red) imaged from 30 hpf.

**Video S2** (related to Figure 1B). Dynamics of junctional materials during the patch-to-ring transition. Time-lapse series with expression of Cdh5-Venus (green) and ZO1-Tdtomato (red) imaged from 30 hpf.

**Video S3** (related to Figure 1C). Recruitment of Rasip1 to the junctional patches. Time-lapse series with expression of Cdh5-Venus (green) and Rasip1-scarlet-I (red) imaged from 30 hpf.

**Video S4** (related to Figure 2I). Degenerations of junctional rings and lumens in *rasip1*$^{ubs28}$ mutants. Time-lapse series with expression of Cdh5-Venus (green) and GFP-Podxl1 (red) imaged from 32 hpf.

**Video S5** (related to Figure 3E’). Photo-conversions on half junctional rings in WT embryos. Time-lapse series with unconverted Cdh5-mClav (green) and converted Cdh5-mClav (red) imaged from 32 hpf.

**Video S6** (related to Figure 3G'). Photo-conversions on half boundary regions of the apical compartments in *rasip1*$^{ubs28}$ mutants. Time-lapse series with unconverted Cdh5-mClav (green) and converted Cdh5-mClav (red) imaged from 32 hpf.

**Video S7** (related to Figure 4A). Cdh5 dynamics in WT embryos. Time-lapse series with expression of Cdh5-Venus imaged from 32 hpf.

**Video S8** (related to Figure 4B). Cdh5 clusters in the apical domains in *rasip1*$^{ubs28}$ mutants. Time-lapse series with expression of Cdh5-Venus imaged from 32 hpf.

**Video S9** (related to Figure 4C). Linear Cdh5 fragments in the apical domains in *rasip1*$^{ubs28}$ mutants. Time-lapse series with expression of Cdh5-Venus imaged from 32 hpf.

**Video S10** (related to Figure S4B). Automatic tracking of Cdh5 clusters in *rasip1*$^{ubs28}$ mutants through PIV. Time-lapse series with expression of Cdh5-Venus imaged from 32 hpf and corresponding quiver plots.

**Video S11** (related to Figure 5A). Dynamics of Myl9a-GFP during the patch-to-ring transition in WT embryos. Time-lapse series with expression of Cdh5-Venus (green) and Myl9a-GFP (red) imaged from 30 hpf.

**Video S12** (related to Figure 5C). Dynamics of Myl9a-GFP during the failed patch-to-ring transition in *rasip1*$^{ubs28}$ mutants. Time-lapse series with expression of Cdh5-Venus (green) and Myl9a-GFP (red) imaged from 30 hpf.

**Video S13** (related to Figure 5G). Dynamics of Myl9a-GFP in the apical compartment and junctions of WT embryos. Time-lapse series with expression of Cdh5-Venus (green) and Myl9a-GFP (red) imaged from 32 hpf.

**Video S14** (related to Figure 5H). Dynamics of Myl9a-GFP in the apical compartment and junctions of *rasip1*$^{ubs28}$ mutants. Time-lapse series with expression of Cdh5-Venus (green) and Myl9a-GFP (red) imaged from 32 hpf.

**Video S15** (related to Figure 6D and 6E). Enrichment of Rasip1 at the constricting junctions and apical domains. Time-lapse series with expression of Cdh5-Venus (green) and Rasip1-scarlet-I (red) imaged from 32 hpf.

**Video S16** (related to Figure 6F). Relocation of Rasip1 from the apical domains to the boundary (junctions). Time-lapse series with expression of GFP-Rasip1 (green) and UCHD-mRuby (red) imaged from 32 hpf.

**Video S17** (related to Figure 6G). Relocation of Rasip1 from boundary to the apical domains. Time-lapse series with expression of GFP-Rasip1 (green) and UCHD-mRuby (red) imaged from 32 hpf.

**Video S18** (related to Figure 6I). Rasip1 clusters colocalized with Myl9a clusters along junctions. Time-lapse series with expression of Myl9a-GFP (green) and Rasip1-scarlet-I (red) imaged from 32 hpf.

**Video S19** (related to Figure 6J). Rasip1 clusters colocalized with Myl9a clusters within the apical compartments. Time-lapse series with expression of Myl9a-GFP (green) and Rasip1-scarlet-I (red) imaged from 32 hpf.

**Video S20** (related to Figure 7H). Activation of opto-RhoA selectively at the apical compartment induced reticulated junctions in WT embryos. Time-lapse series with expression of Cdh5-Venus (green) and RhoA-BcLOV4-mCherry (red) imaged from 32 hpf. Blue labels the ROI of activation.
